# A single-cell atlas of human fetal lung development between 14 and 19 weeks of gestation

**DOI:** 10.1101/2021.12.23.473945

**Authors:** Laurent Renesme, Flore Lesage, David Cook, Shumei Zhong, Satu Hänninen, Olli Carpén, Ivana Mižíková, Bernard Thébaud

## Abstract

**Rationale:** Human lung development has been mainly described in morphologic studies and the potential underlying molecular mechanisms were extrapolated from animal models. Therefore, there is a need to gather knowledge from native human lung tissue. In this study we describe changes at a single-cell level in human fetal lungs during the pseudoglandular stage.

**Methods:** We report the cellular composition, cell trajectories and cell-to-cell communication in developing human lungs with single-nuclei RNA sequencing (snRNA-seq) on 23,251 nuclei isolated from nine human fetuses with gestational ages between 14 to 19 weeks of gestation.

**Results:** We identified nine different cell types, including a rare pulmonary neuroendocrine cells population. For each cell type, marker genes are reported, and selected marker genes are used for spatial validation with fluorescent RNA *in situ* hybridization. Enrichment and developmental trajectory analysis provide insight into molecular mechanisms and signaling pathways within individual cell clusters according to gestational age. Lastly, ligand-receptor analysis highlights determinants of cell-to-cell communication among the different cell types through the pseudoglandular stage, including general developmental pathways (NOTCH and TGFB), as well as more specific pathways involved in vasculogenesis, neurogenesis, and immune system regulation.

**Conclusion:** These findings provide a clinically relevant background for research hypotheses generation in projects studying normal or impaired lung development and help to develop and validate surrogate models to study human lung development, such as human lung organoids.

**TAKE HOME MESSAGE:** Using a single-cell transcriptomic approach (single-nuclei RNA sequencing), we describe here, for the first time, the cellular landscape, cell developmental trajectories, and cell-to-cell communication in the developing human lung during the pseudoglandular stage.

## INTRODUCTION

Human lung development is traditionally divided into 5 stages: embryonic (1-7 weeks of gestation (WG)), pseudoglandular (5-17 WG), canalicular (16-26 WG), saccular (24 to 38 WG), and alveolar (36 WG to 3 years, with late alveolarization continuing up to early adulthood)^1^. While these stages have been defined morphologically and histologically during human fetal development, very little is known about the underlying molecular and cellular mechanisms^2,3^. Detailed insight into the molecular pathways and cellular communication which contribute to lung development would improve our understanding of lung repair as it may recapitulate fetal lung development, and would enable us to develop new therapeutic targets^4,5^. Due to developmental similarities in human and rodent lungs, mouse and rats have been used extensively to study lung development^6^. Airway branching morphogenesis in rodents is well described, and gain- and loss-of-function studies using genetically modified animals have identified the critical factors of this developmental process^3,7^. However, some compartments of the lung, such as stroma or vascular and neuronal networks, remain poorly explored during the fetal developmental period despite their potential key roles in lung development^8,9^. While versatile in experimental settings, rodent models of lung development have several limitations, including species-specific maturation and function at birth, and molecular regulation^10,11^. Therefore, there is a need to unravel cellular composition, developmental trajectories, and signaling within the developing human lung.

Single-cell and single-nuclei RNA sequencing (scRNA-seq and snRNA-seq, respectively) can reveal complex and rare populations, uncover regulatory relationships, and track distinct cell lineages in development. In addition, computational methods have been developed to analyze cellular communications and to determine which extracellular signals influence the cell population of interest.

In the present study, we report the cellular composition, developmental pathways and cell-to-cell communication in fetal human developing lungs using snRNA-seq approach on 23,251 nuclei isolated from nine lungs with gestational ages (GAs, described in WG) spanning through the pseudoglandular and canalicular stages (14 to 19 WG). We describe nine different cells populations, including rare cell population of pulmonary neuroendocrine cells (PNEC). For each identified population we report marker genes and enrichment and developmental state analysis. Additionally, we provide a spatial validation for selected populations using fluorescent RNA *in situ* hybridization (FISH). Lastly, ligand-receptor analysis highlights determinants of cell-to-cell communication among the different cell types.

## METHODS

A full description of the experimental procedures can be found in the supplementary methods.

### Human fetal tissue collection

De-identified human fetal lung samples were obtained under REB approval by The Ottawa Hospital Review Ethical Board (20170603-01H). Lung tissue was snap frozen and stored at -80°C.

### Single-nuclei suspension preparation

Single nuclei isolation was performed according to Martelotto et al.^12^ with minor adjustments. Sample multiplexing was performed using the MULTI-seq protocol as reported before^13,14^. Nuclei integrity was confirmed by fluorescent microscopy (Axio Imager M2, Carl Zeiss, Toronto, ON, Canada) and single diploid nuclei were sorted using a flow cytometer (BD LSR Fortessa, Beckton Dickinson Biosciences, Franklin Lakes, NJ, USA) (Supplemental figure 1a-b)

### Single-nuclei sequencing

Cell capture and library production was performed with the Chromium system (10X Genomics, Pleasanton, CA, USA). Sequencing was performed with NextSeq500 (Illumina, San Diego, CA, USA). Raw sequencing reads were processed using CellRanger v3.1.0. All analyses were performed with Seurat v4.0.0^15^. SCTransform() was used to normalize samples, select highly variable genes, and to regress out cell cycle and cell stress effects. To eliminate batch effects or biological variability effects on clustering, the data integration method implemented by Seurat for SCTransform-normalized data was performed, using the SelectIntegrationFeatures(), PrepSCTIntegration(), FindIntegrationAnchors(), and IntegrateData() functions. Differential State Analysis (DSA) was performed using muscat R package^16^. Gene set enrichment analysis (GSEA) was performed with Metascape^17^. Cell communications were inferred using NicheNet (v1.0.0) R package^18^.

### In situ hybridization

FISH was performed on formalin-fixed, paraffin embedded human fetal lungs. 3μm thick sections were analysed using the RNAscope technology following the manufacturer’s protocol, and as previously described^14^.

## RESULTS

### Cellular landscape of the human developing lung

To characterize the cellular composition in developing human lungs, we generated snRNA-seq profiles of 23,251 nuclei isolated from nine fetal lung samples with GAs between 14+1 and 19+0 weeks (Figure 1a, and Supplemental figure 1c). We identified nine distinct cell populations, present across all GAs (Figure 1b-d), and described the lung cellular composition variation over time. While proportions of both stromal populations decreased over time (83.7 and 6.1% at 14+1 weeks *vs*. 47.6 and 4% at 19+0 weeks), the size of distal airway epithelium cluster increased (from 3.6% to 39.5%) (Figure 1c and Supplemental table 1). Each cell cluster displayed a unique molecular signature characterized by multiple DEGs (Figure 1e and Supplemental table 2), and cell populations were annotated based on the expression of canonical cell lineage markers (Figure 1f and 2a). Additionally, among the top ten DEGs in some clusters, we identified less commonly used transcriptional patterns (Figure 1e). For example, pericytes expressed genes involved in the cGMP-PKG pathway such as *MEF2C, PDE3A, TRPC6*, and *PLCB*. Vascular and lymphatic endothelium expressed genes involved in angiogenesis, respectively *EPAS1, PTPRB, CALCRL* and *NRP2, VAV3, STAB2*. Finally, the immune cells cluster expressed *HDAC9, SAMSN1* and *DOCK8* that are involved in lymphocyte activation.

**Figure 1:**
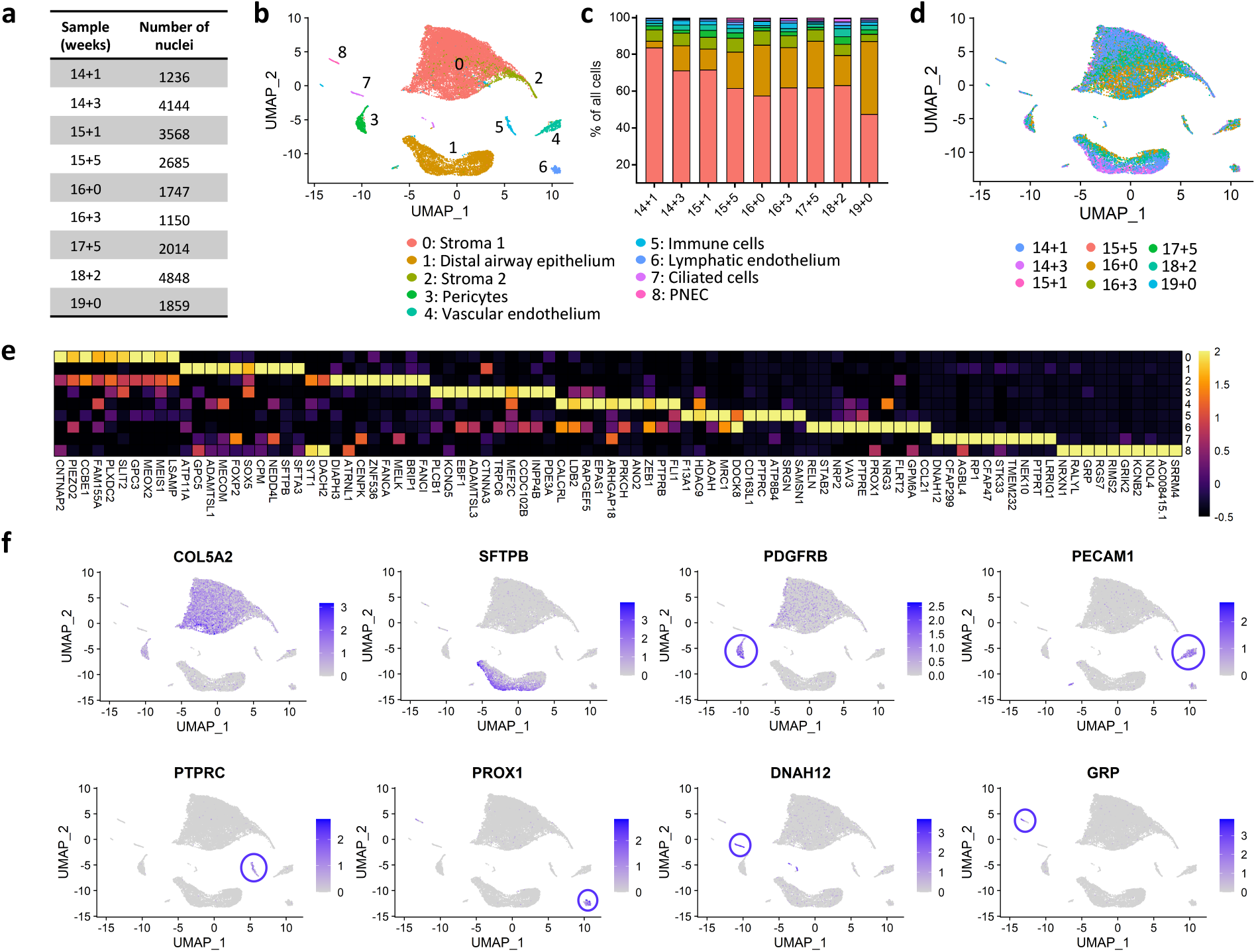
Cellular composition of human fetal lung tissue between 14 and 19 weeks of gestation. a) Table depicting the number of nuclei sequenced in each sample. b) UMAP plot depicting 9 cell clusters identified by snRNA-seq analysis of fetal lung tissue collected between 14 and 19 weeks of gestation. c) Bar graph depicting the percentual contribution of individual cell populations at different developmental stages. d) UMAP plot depicting the distribution of fetal lung cells based on developmental age. e) Heatmap of top ten most differentially expressed genes across fetal lung cell clusters depicted in panel b. f) UMAP plots showing expression levels for canonical markers of lung cell populations, including stroma (COL5A2), distal airway epithelium (SFTPB), pericytes (PDGFRB), vascular endothelium (PECAM1), immune cells (PTPRC), lymphatic endothelium (PROX1), ciliated cells (DNAH12) and pulmonary neuroendocrine cells (GRP). The intensity of expression is indicated by purple coloring. Expression values in heatmaps and UMAP plots represent Z-score-transformed log(TP10k+1) values. Log(TP10k+1) corresponds to log-transformed UMIs per 10k.

To further confirm clusters identities, and to gain further insights into the underlying molecular functions and biological processes, we performed an GSEA. Cell-type specific terms were associated with all the identified clusters. As expected, stromal populations 1 and 2 were associated with general terms such as extracellular matrix and mesenchyme development. Distal airway epithelium was associated with respiratory gaseous exchange, as well as apical and basolateral plasma membrane terms, while ciliated cells were characterized by cilium-specific pathways. Blood vessel development was associated with both, pericytes and vascular endothelium, while lymph vessel development pathways were specific to lymphatic endothelium cluster. Immune cells cluster was characterized by interleukin signalling and myeloid cell differentiation pathways. Finally, transsynaptic signaling was associated specifically with pulmonary neuroendocrine cells (Figure 2b, Supplemental table 3).

**Figure 2:**
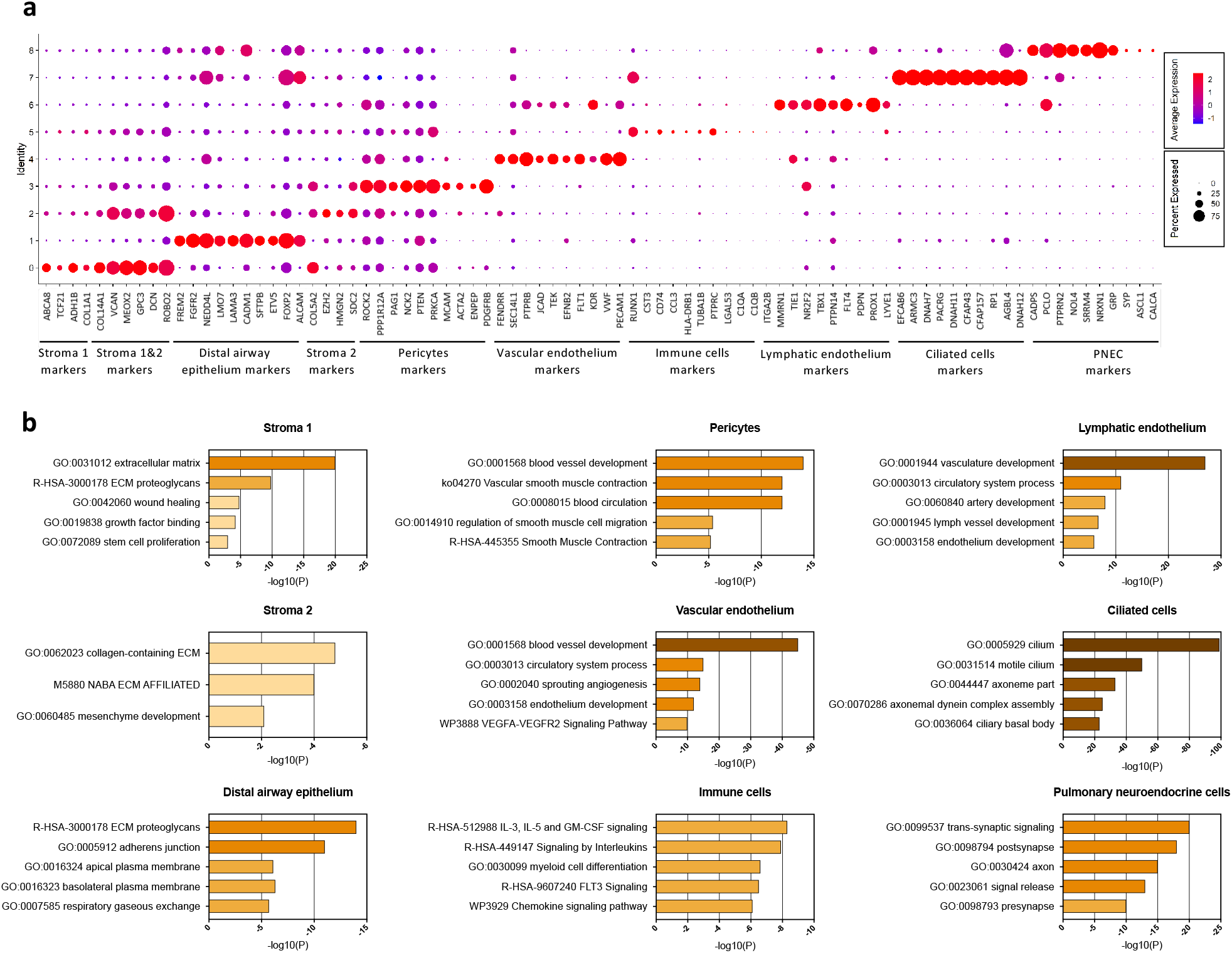
Identification of cell clusters comprising human fetal lung between 14 and 19 weeks of gestation. a) Dotplot depicting the expression of most commonly used markers for each cell cluster as described in literature. Expression levels in the dotplot are presented as log(TP10k+1) values. Log(TP10k+1) corresponds to log-transformed UMIs per 10k. b) Selected regulated pathways for each cluster identified by gene set enrichment analysis. All terms are significantly enriched (adjusted p-value < 0.05). Plotted values represent -log10(p-value).

### Developmental changes in gene expression in developing lung stromal cells

Within the stromal population (stroma 1 and 2), we identified three subclusters with distinct gene expression patterns represented across different GAs (Figure 3a-e, Supplemental table 4). Stromal subclusters 0 and 1 expressed traditional matrix fibroblast markers (*FN1, MEOX2*) (Figure 3f). Pulmonary fibroblasts can be typically segregated into two subtypes based on the distinct expression of *COL13A1* and *COL14A1*^19^. In our dataset the expression of *COL13A1* was most prominent in stromal subcluster 0, which was also characterized by expression of additional genes associated with *COL13A1*^+^ fibroblasts (*ITGA8, LIMCH1, MYLK, PLXDC2, MACF1*) (Figure 3f). A sizeable fraction of subcluster 1 expressed *COL14A1*, as well as other matrix fibroblast markers (*FBLN1, COL1A2, AKAP12, COL1A1*) (Figure 3f). Stromal subcluster 2 showed similarities to mesenchymal progenitors as described by *Xie et al*.^19^ and the LungGENS database^20^, characterized by the expression of *SDC2* and *SMARCC1* (Figure 3f). However, this sub-population did not express any proliferative genes such as *TOP2A* or *MKI67* which are typically expressed in self-renewing progenitor populations. GSEA of stromal subclusters revealed supportive roles for the stromal cells in lung and mesenchyme development, with subcluster 0 and 1 being associated with regulation of organ architecture (connective tissue development, cell junction assembly, cytoskeleton organization, negative regulation of locomotion/cell motility) and subcluster 2 being associated with proliferative/stemness functions (DNA binding transcription factor activity, negative regulation of cell differentiation) (Supplemental figure 2a and c, Supplemental table 5).

**Figure 3.**
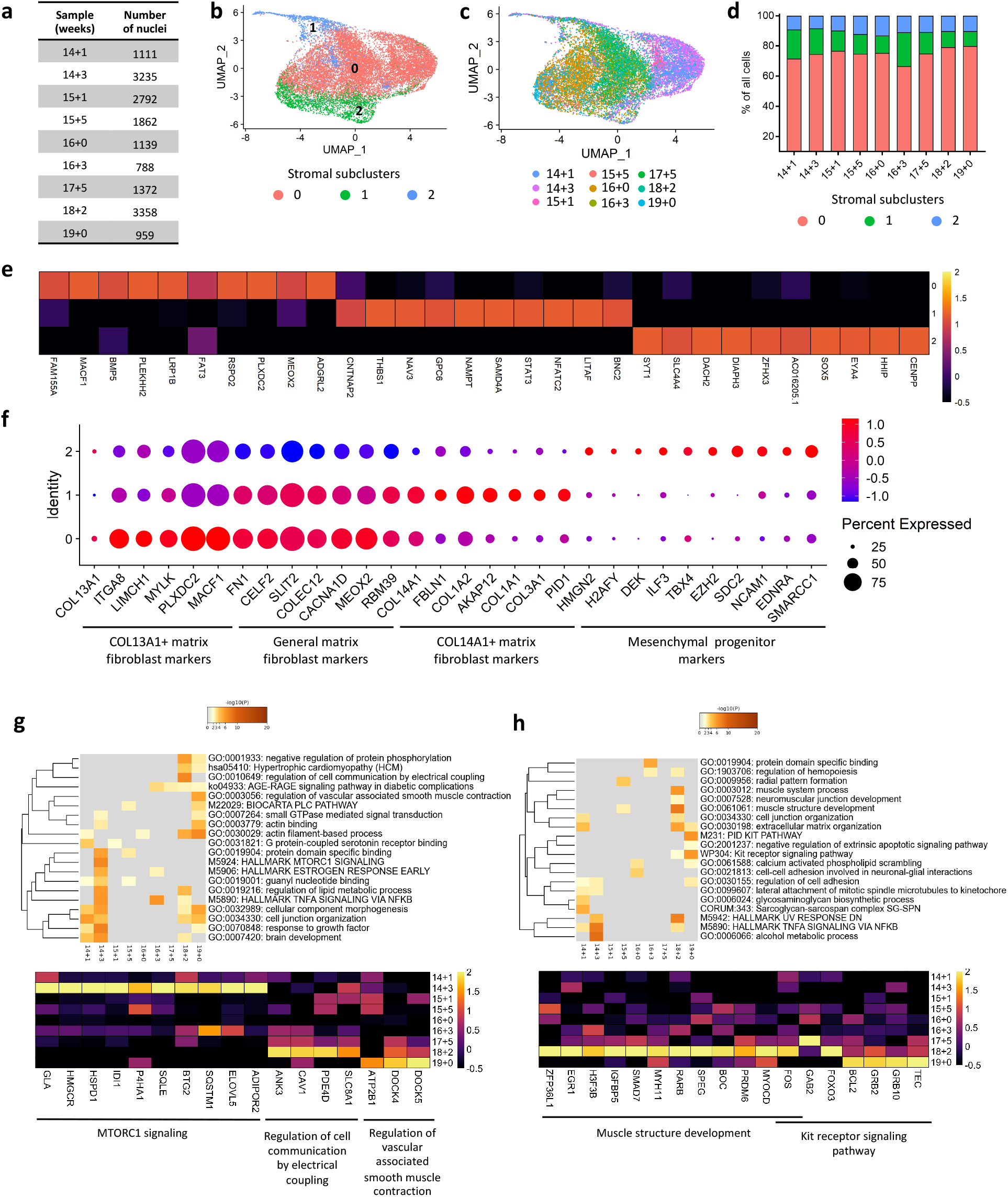
Cellular composition and developmental changes in gene expression of stromal clusters in human fetal lung between 14 and 19 weeks of gestation. a) Table depicting the number of stromal cells nuclei sequenced in each sample. b) UMAP plot depicting the 3 stromal subclusters identified within the fetal lung stroma. c) UMAP plot depicting the distribution of fetal lung stromal cells based on the developmental age. d) Percentual contribution of the different lung stromal subclusters in each individual sample. e) Heatmap depicting the top ten most differentially expressed genes across fetal lung stromal subclusters shown in panel b). f) Dotplot depicting the expression of stromal markers known to be associated with *COL13A1*^+^ matrix fibroblasts (subcluster 0), *COL14A1*^+^ matrix fibroblasts (subcluster 1), general matrix fibroblasts (subcluster 0 and 1), and mesenchymal progenitors (subcluster 2) in stromal subclusters. g) Heatmaps depicting the enriched terms associated with individual GAs as identified by multi-list enrichment analysis based on DSA in the main cluster stroma 1 (top), and heatmap depicting the expression level of genes associated with selected enriched terms (bottom). h) Heatmaps depicting the enriched terms associated with individual GAs as identified by multi-list enrichment analysis based on DSA in the main cluster stroma 2 (top), and heatmap depicting the expression level of genes associated with selected enriched terms (bottom). Expression levels in the heatmaps and dotplots are presented as log(TP10k+1) values. Log(TP10k+1) corresponds to log-transformed UMIs per 10k.

Next, we performed DSA-based GSEA (Supplemental tables 6, 7 and 8). In the two main stromal clusters, Stroma 1 cluster was associated with response to growth factor and MTORC1 signaling for early GA (14+1 to 14+3 WG), and regulation of cell communication by electrical coupling, regulation of vascular associated smooth muscle contraction for late GA (18+2 to 19+0 WG) (Figure 3g, Supplemental figure 3). For stroma 2, prominent regulated enriched pathways included TNFα signaling via NFκB for early GA, as well as muscle structure development and Kit receptor signaling pathway for late GA (Figure 3h, Supplemental figure 3).

### Developmental changes in gene expression in developing airway epithelium

Upon subclustering of the distal airway epithelium, we identified three subclusters with distinct gene expression represented across the different GAs (Figure 4a-e, Supplemental table 4). Epithelial subclusters shared common marker genes with bud tip progenitors (subcluster 0: *SFTPC, ETV5*), bud tip adjacent (subcluster 1: *DMD, MECOM*), and secretory progenitor (subcluster 2: *SCGB3A2, CFTR*) epithelial cells as identified by *Miller et al*.^21^ in human fetal lung tissue of similar GA (Figure 4f). In addition, a GSEA was performed with DEG from these three subclusters (Supplemental figure 2b and d, Supplemental table 5). Based on DEG and DSA results, we selected gene markers for spatial validation on the fetal lung tissue by FISH (Supplemental tables 2 and 6). Due to their cluster specificity, we selected *SEMA3C* and *SFTPB* as potential early and late distal airway epithelium markers in fetal lung (Supplemental figure 4a). As seen by FISH (Figure 4g), distal airway epithelium (marked by *FGFR2*^+^) in earlier GAs co-expresses *SEMA3C*, with only few cells also co-expressing *SFTPB*. However, the majority of *FGFR2*^+^ epithelial cells surrounding the airway lumen at later GAs were *SEMA3C*^-^ and *SFTPB*^+^. Notably, an adjacent layer of *FGF2*^*-*^/*SEMA3C*^+^/*SFTPB*^-^ cells can also be seen (Figure 4h). Finally, based on the DSA, we performed a GSEA. Among the most regulated pathways in distal airway epithelium were sensory organ development and blood vessel development (Figure 4h, Supplemental figure 3, Supplemental tables 7 and 8).

**Figure 4.**
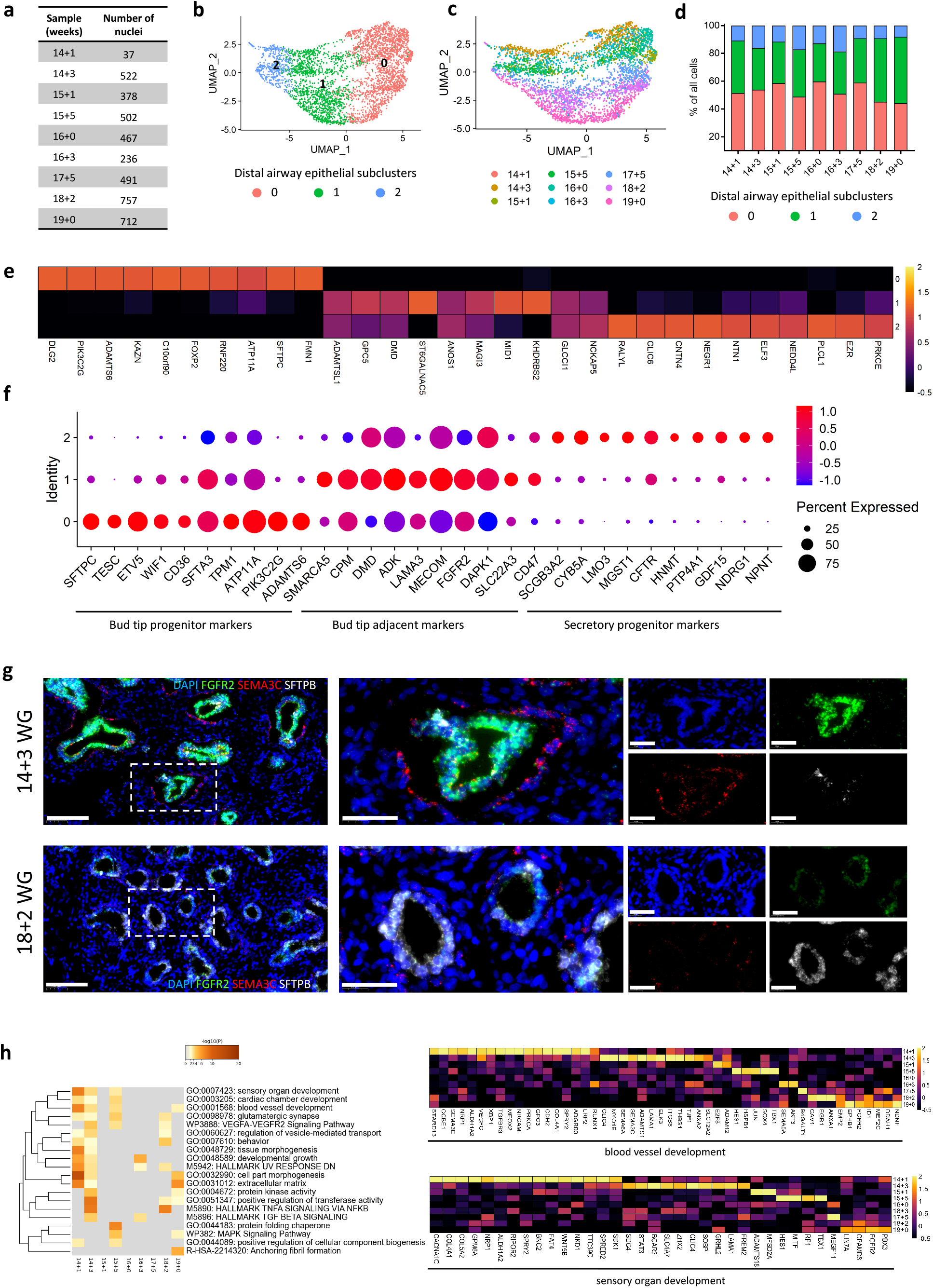
Cellular composition, developmental changes in gene expression, and spatial localization of distal airway epithelial cells in human fetal lung between 14 and 19 weeks of gestation. a) Table depicting the number of distal airway epithelial cells nuclei sequenced per sample. b) UMAP plot depicting the 3 epithelial subclusters identified within the distal airway epithelium cluster; c) UMAP plot depicting the distribution of fetal lung distal airway epithelium cells based on developmental age. d) Percentage contribution to the different lung epithelial subclusters per sample. e) Heatmap depicting the top ten most differentially expressed genes across fetal lung epithelial subclusters shown in panel b). f) Dotplot depicting the expression of epithelial markers known to be associated with bud tip progenitor cells (subcluster 0), bud tip adjacent cells (subcluster 1) and secretory progenitors (subcluster 2). g) Representative fluorescent RNA *in situ* hybridization pictures showing co-expression of *FGFR2* (green), *SEMA3C* (red) and *SFTPB* (white) in the airway epithelium in human fetal lungs at 14+3 (top) and 18+2 (bottom) weeks of gestation. Magnification at x20 (scale bar 100 µm) and x63 (scale bar 40 µm). Expression levels in the heatmaps and dotplots are presented as log(TP10k+1) values. Log(TP10k+1) corresponds to log-transformed UMIs per 10k. h) Heatmaps depicting the enriched terms associated with individual GAs as identified by multi-list enrichment analysis based on DSA in the main cluster distal airway epithelium (left), and heatmap depicting the expression level of genes associated with selected enriched terms (right).

In the case of ciliated cells and PNECs, DSA identified only few DEGs, confirming previous reports on the early specification of these cell populations (Supplementary figures 5 and 6, respectively)^22,23^. Interestingly, only a very small fraction of the PNEC population showed high expression of typical PNEC markers such as *GRP1, CALCA, ASCL1*, and *SYP*^21^, whereas markers associated with the synapse, signal release and transsynaptic signaling (*NRXN1, PTPRN2, PCLO, CADPS*) were expressed in the majority of the PNECs in our dataset. FISH was performed using *NRXN1* and *GRP* as PNEC markers (Supplemental figure 4b). The *NRXN1*^+^/*GRP*^+^ PNECs were present at all GAs and were either adjacent to or localized within the airway epithelium. Both, isolated cells and small cells clusters (neuroendocrine bodies) were noted. Interestingly, while all *GRP*^+^ cells were *NRXN1*^+^, *GRP*^-^/*NRXN1*^+^ cells were also noted, suggesting a possible existence of two subpopulations of PNECs in the developing lung (Figure 5).

**Figure 5.**
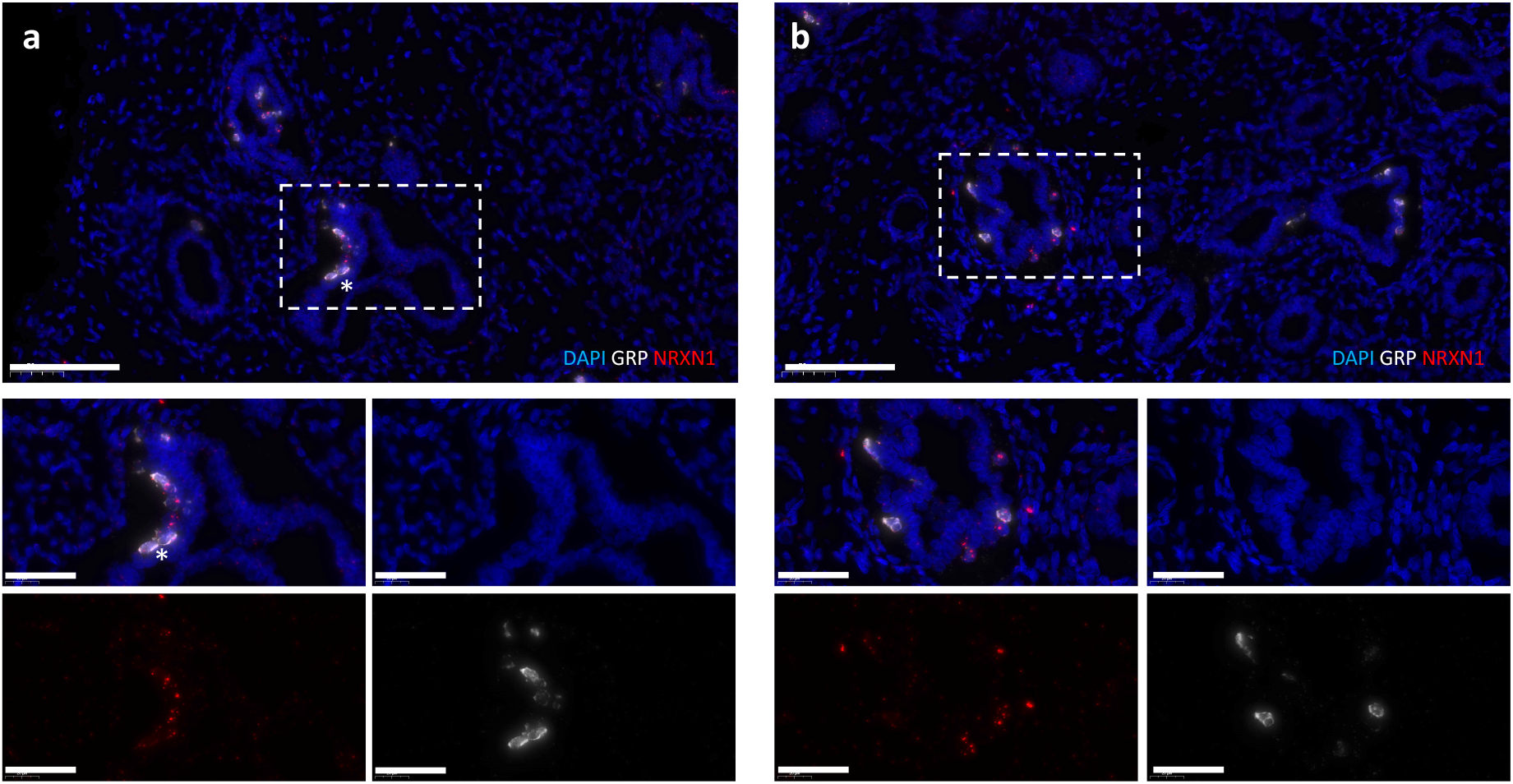
Fluorescent RNA in situ hybridization of pulmonary neuroendocrine cells in the human fetal lung. Representative fluorescent RNA *in situ* hybridization pictures showing co-expression of *NRXN1* (red) and *GRP* (white) in PNEC in human fetal lungs at a) 14+3 and b) 15+5 weeks of gestation. Both, isolated *NRXN1*^+^/*GRP*^+^ PNECs and small PNECs clusters (neuroendocrine bodies, white *) were noted. Magnification at x20 (top, scale bar 100 µm) and x63 (bottom, scale bar 40 µm).

**Figure 6.**
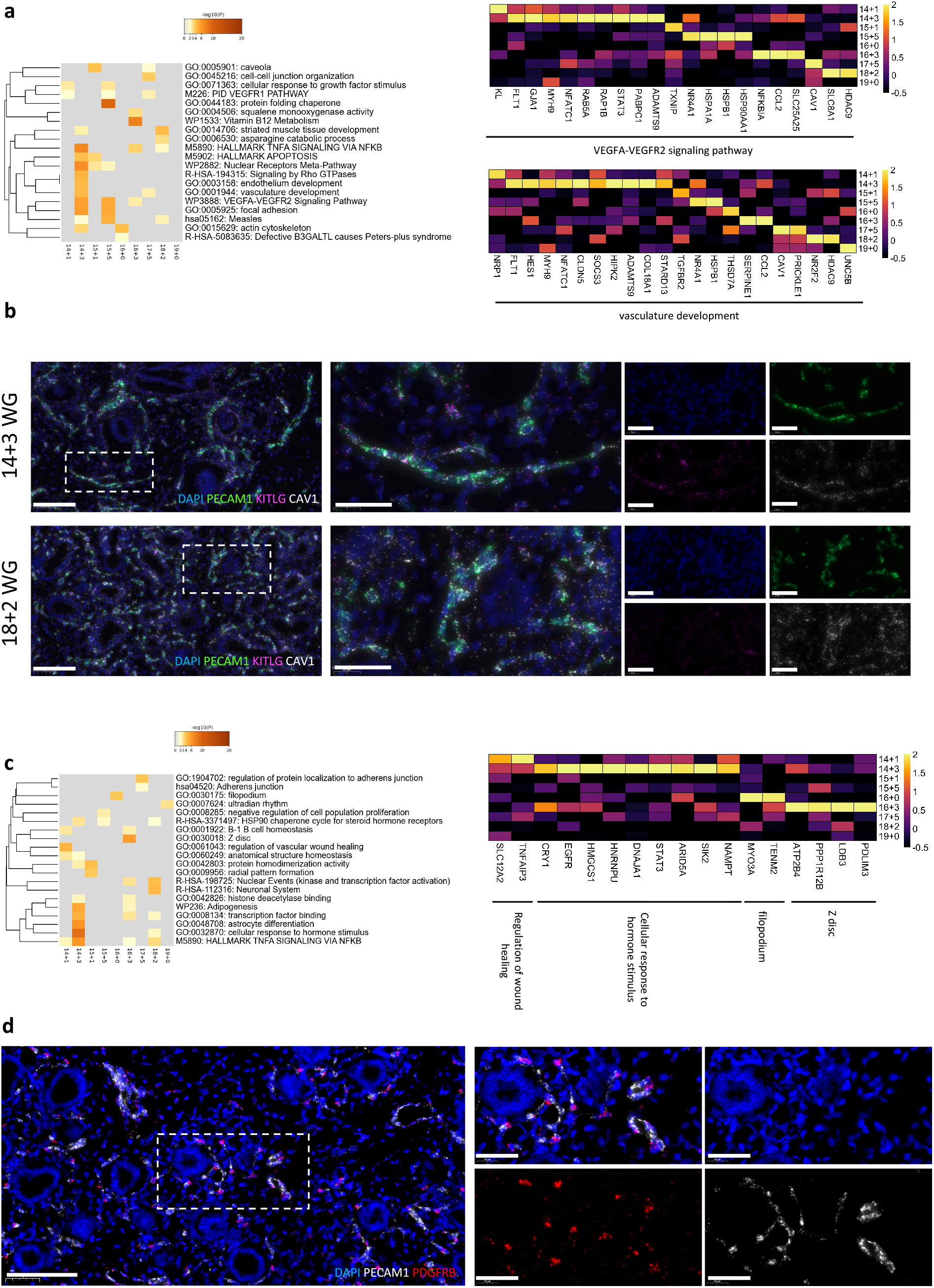
Developmental changes in gene expression and spatial localization of vascular endothelium and pericytes. a) Heatmaps depicting the enriched terms associated with individual GAs as identified by multi-list enrichment analysis based on DSA in the main cluster vascular endothelium (left), and heatmap depicting the expression level of genes associated with selected enriched terms (right). b) Representative fluorescent RNA *in situ* hybridization pictures showing co-expression of *PECAM1* (green), *KITLG* (magenta) and *CAV1* (white) in the vascular endothelium in human fetal lungs at 14+3 (top) and 18+2 (bottom) weeks of gestation. Magnification at x20 (scale bar 100 µm) and x63 (scale bar 40 µm). c) Heatmaps depicting the enriched terms associated with individual GAs as identified by multi-list enrichment analysis based on DSA in the main cluster pericytes (left), and heatmap depicting the expression level of genes associated with selected enriched terms (right). Expression levels in the heatmaps are presented as log(TP10k+1) values. Log(TP10k+1) corresponds to log-transformed UMIs per 10k. d) Representative fluorescent RNA *in situ* hybridization pictures showing co-localization of *PECAM1+* endothelial cells (white) and *PDGFRB+* pericytes (red) in human fetal lungs at 18+2 weeks of gestation. Magnification at x20 (scale bar 100 µm) and x63 (scale bar 40 µm).

### Developmental changes in gene expression in developing endothelial, pericyte and immune cells populations

Vascular endothelium at early GA (14+3 WG) was characterized by cell type specific pathways such as VEGFA-VEGFR2 signaling pathway, endothelium development and vasculature development (Figure 6a, Supplemental figure 3). In the lymphatic endothelium, genes expressed at early GAs were involved in ubiquitin protein ligase binding and IL-6 signaling. Interestingly, ubiquitination plays a central role in NOTCH signaling^24^ and IL-6 is involved in lymph angiogenesis via VEGF-C^25^, whereas later GAs were characterized by expression of genes involved in “cholesterol biosynthesis” (Supplemental figure 7). Next, to visualize the vascular endothelial cells in the fetal lung we performed a FISH (Figure 6b). Based on the DSA (Supplemental table 6), we selected KITLG as an early, and CAV1 as a late marker for vascular endothelium during the pseudoglandular phase (Supplemental figure 4c). However, both KITLG^+^/PECAM^+^ and CAV1^+^/PECAM^+^ double positive cells could be found at all investigated GAs (Figure 6c).

The pericytes cluster showed gene expression associated with regulation of vascular wound healing and cellular response to hormone stimulus at early GAs. In addition, cellular components involved in pericytes functions, such as filipodium and Z disc were described at later GAs (16+0 and 16+3 WG, respectively)^26,27^ (Figure 6c). *PDGFRB*^+^ pericytes (Supplemental figure 4d) could be found in immediate proximity to vascular endothelium (defined by the expression of *PECAM1*^+^) in the lungs throughout the pseudoglandular development (Figure 6d).

Finally, in the case of immune cells population, the DEGs at early GAs were associated with “lymphocyte activation” and “TNFα signaling via NFκβ” which is known to be crucial in lymphocyte activation, as well the specification of innate and adaptive immunity^28^ (Supplemental figure 8).

### Cellular crosstalk in the human developing lung

In order to understand cell communication networks within the developing human fetal lung samples, we performed a ligand-receptor interaction analysis using NicheNet^18^. The analysis focused on time-dependent changes in cell communication rather than stable cell crosstalk, and the set of gene of interest used for the analysis was defined by differential expression between early GAs (14+1 and 14+3 WG, reference condition) and late GAs (18+0 and 19+0 WG, condition of interest). Cellular crosstalks identified at later GAs are presented in Figure 7, with the identified receptors and predicted target genes involved (Figure 8). GSEA on predicted target genes are presented in Supplemental table 9. Inferred cellular communications at later GAs were associated with general pathways such as NOTCH and TGFB pathways, as well as in pathways involved in immune system regulation, and neurogenic tissue and vasculature development.

**Figure 7:**
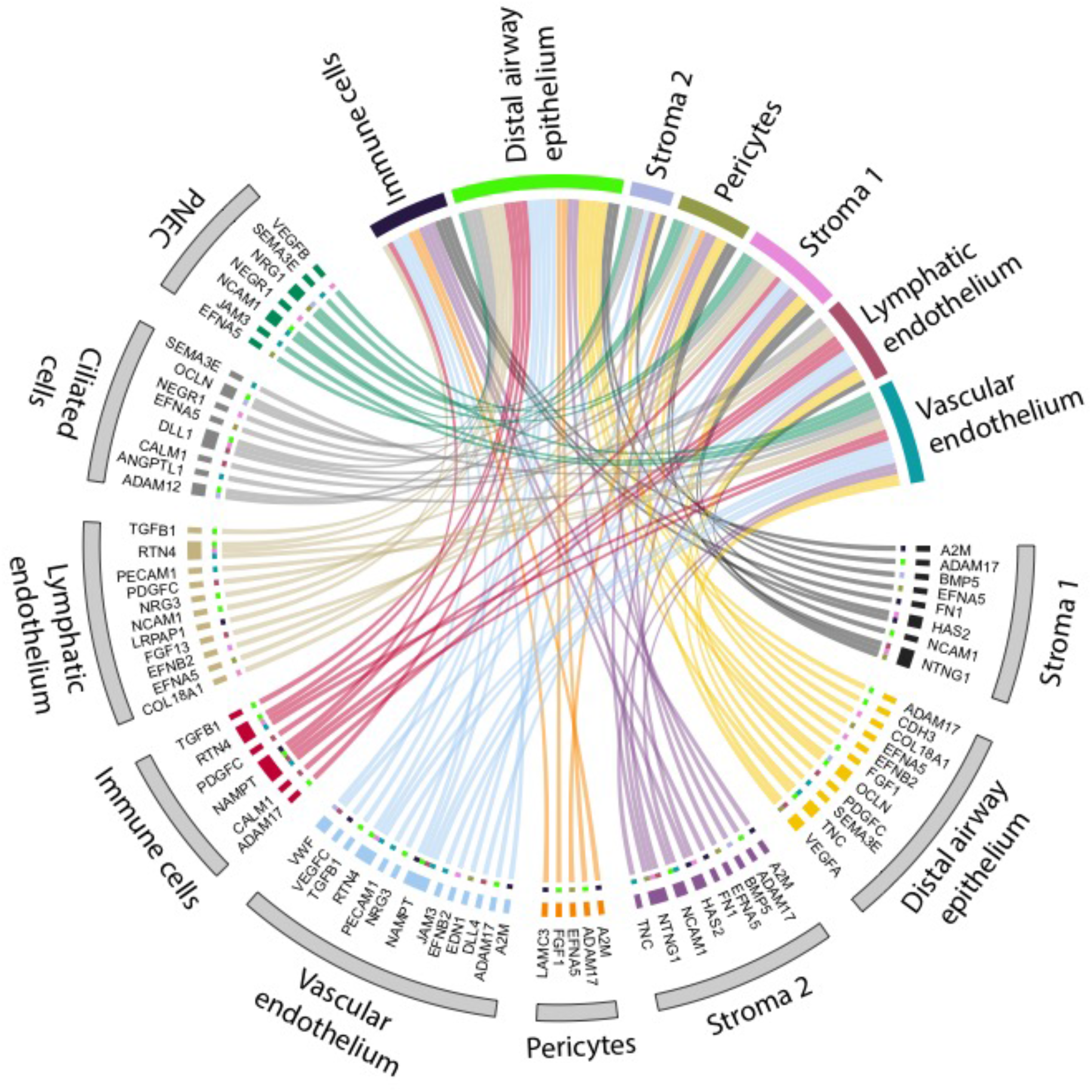
Cell communication pathways induced over time. Circos plot showing inferred cell communications identified with NicheNet by comparing the two latest GA (18+2 and 19+0 weeks) with the two earliest GA (14+1 and 14+3 weeks). Colored populations in the upper part of the plot represent the receiving cell populations, while grey-colored populations in the lower part of the plot represent the signal senders and their ligands. Each ligand is connected to the respective receiving population.

**Figure 8.**
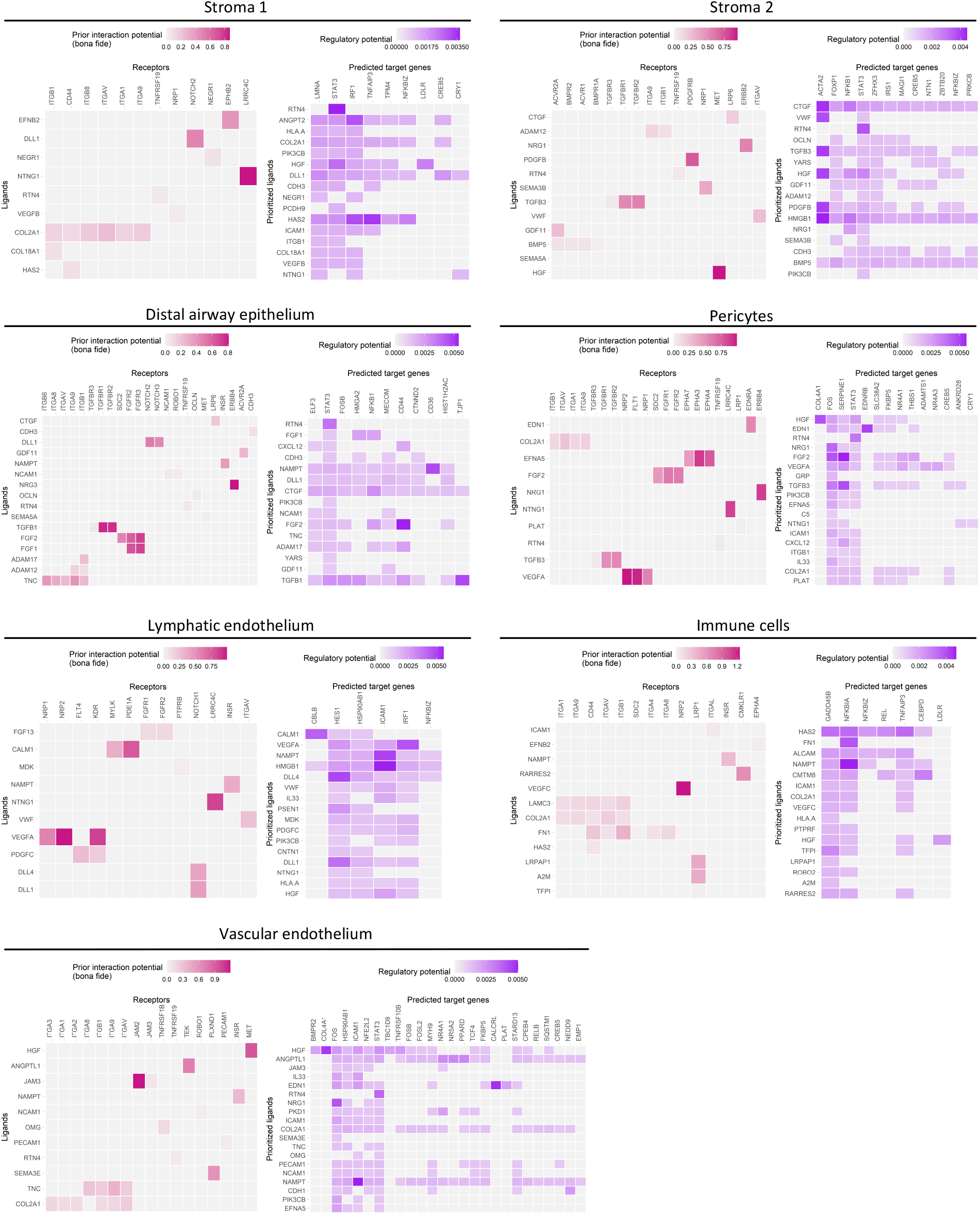
Identified receptors and predicted target genes involved in cell communication. Heatmaps showing bona-fide receptors and predicted target genes for each receiving cell population identified with NicheNet by comparing the two latest GA (18+2 and 19+0 weeks) with the two earliest GA (14+1 and 14+3 weeks). Receiving cell populations include stroma 1 and 2, pericytes, Lymphatic endothelium, distal airway epithelium, immune cells and vascular endothelium. For each cell population, pink heatmap reports bona fide receptors and their ligands and purple heatmap shows predicted target genes according to prioritized ligands.

We identified key factors in the NOTCH signaling pathway such as *DLL1* (expressed by the ciliated cells) and its receptors *NOTCH2* (expressed in stroma 1 and distal airway epithelium) and *NOTCH3* (distal airway epithelium). Predicted target genes were associated with T cell differentiation and TNF signaling in stroma 1 cluster, and with cellular senescence in airway epithelium. Moreover, *DLL4* expressed by the vascular endothelium targeted changes in gene expression associated with INFγ-mediated signaling and leukocytes cell-cell adhesion in lymphatic endothelium via its receptor *NOTCH1. ADAM17*, an important NOTCH signaling activator, was expressed by majority of cell populations, while its receptor *ITGB1* was expressed by distal airway epithelium. Potential *ADAM17* target genes were primarily associated with inflammatory response.

*TGFB1* was expressed by lymphatic and vascular endothelium and the immune cells. Its receptors, *TGFBR1* and *TGFBR2*, were expressed in distal airway epithelium and the predicted target genes were associated with cellular senescence and gastrin signaling. *BMP5*, another member of the TFGB family, was expressed in both stromal clusters, while its receptors (*BMPR2, BMPR1A, ACVR2A* and *ACVR1*) were specifically expressed by stroma 2. The targeted genes were associated with insulin resistance, T cell differentiation, IL-18 signaling, and with regulation of cell adhesion and motility.

*CD44* was expressed in stroma 1 and immune cells and its potential ligands HAS2 and FN1 were expressed by stroma 1 and 2 clusters. Predicted target genes for CD44 activation in stroma 1 were associated with T cell differentiation and regulation of adaptative immune response. Target genes in immune cells cluster were associated with IL-18 and NF-kappa B signaling pathways.

Further regulated were also genes involved in neurogenic tissue development, including stroma-expressed *NTNG1*, and its receptor *LRRC4C, NEGR1* (expressed by ciliated cells and PNECs) and its receptor NEGR1 (stroma 1), as well as *NCAM1* (stroma, lymphatic, and PNEC clusters) and its receptor *ROBO1* (distal airway epithelium and vascular endothelium). TNC, an extracellular matrix protein implicated in the guidance of migrating neurons, was expressed in stroma 2 and distal airway epithelium clusters, with its receptors expressed in distal airway epithelium and vascular endothelium.

Lastly, in regard to vasculature development, VEGF ligands were expressed by distal airway epithelium (*VEGFA*), vascular endothelium (*VEGFC*), and PNEC (*VEGFB*). *VEGFA* receptors were expressed in pericytes (*NRP1, NRP2, FLT1*) with predicted target genes associated with TGF-beta receptor signaling and positive regulation of cell migration, and in lymphatic endothelium (*NRP1, NRP2, KDR*) with predicted target genes associated with INFγ-mediated signaling and regulation of leukocytes cell-cell adhesion. *VEGFB* receptor *NRP1* was expressed in stroma 1, while *NRP2* was expressed in immune cells, and the predicted target genes were associated with NF-kappa B signaling. In addition, we reported the expression of *SEMA3E* and its receptor *PLXND1*, which are required for normal vascular patterning during embryogenesis. *SEMA3E* was expressed by the airway epithelium (ciliated cells, PNEC and distal airway epithelium) and *PLXND1*, was expressed by the vascular endothelium.

## DISCUSSION

This is the first report of an unbiased transcriptomic analysis of human fetal lung samples, providing a detailed cell atlas of the human developing lung during pseudoglandular and early canalicular stages at a single-cell resolution. We identified 9 cell types, including rare cell type PNEC, and described developmental changes in gene expression and cell-cell communication across GAs. In addition to canonical marker genes, we identified an array of marker genes for each cell population and validated the spatial localization of specific cell populations such as PNEC and pericytes in the human developing lung. Lastly, the cell communication analysis provided unique insight into detailed cellular crosstalk and main pathways involved in the pseudoglandular stage of lung development.

Only few studies to date have reported the use of single-cell transcriptomic on human developing lung, typically including a small number of samples or focusing on specific cell types ^21,29,30^. Our study reported transcriptomics data from 9 fetuses, representing each developmental week from 14 and 19 WG, allowing detailed analyses of gene expression changes overtime. Additionally, choosing single-nuclei isolation from frozen material, rather than single-cell isolation from fresh tissue, enabled us to isolate a relatively large number of distal airway epithelial cells (up to 39.5% of all cells at 19+0 WG). This allowed us to show a steady increase in distal airway epithelial cells across GAs in relation to airway development, whereas previous studies using single-cell isolation approach seemed more prone to dissociation bias (with epithelial cells representing less than 4% of the total cells)^29,30^. These findings suggest that transcriptomic analysis of tissue with high stromal content, such as human fetal lungs, might be more efficient when using a single-nuclei, rather than a single-cell isolation preparation.

The main strength of this study lies within the detailed description of the cellular crosstalk changes between early (14+1 and 14+3 WG) and late (18+2 and 19+0 WG) GAs. We identified several critical pathways associated with normal lung development and lung disease. For instance, VEGF plays a central role in lung development and maintenance, and is involved in the pathogenesis of many lung diseases, including emphysema, chronic obstructive pulmonary disease, bronchopulmonary dysplasia, pulmonary hypertension, acute lung injury, and asthma^31^. ADAM17, a NOTCH activator, plays a key role in lung inflammation regulation by increasing epithelium and smooth muscle cells permeability, secretion of inflammatory mediators, and trans-endothelial leukocyte migration, and its deficiency leads to decreased airway branching ^32^In addition, the role of the neurogenic tissue and vasculature in the developing lung is rapidly gaining interest^8,9^. Here we have identified key factors for neurogenic tissue development also known to be involved in abnormal lung development, such as *ROBO1* (expressed in distal airway epithelium and vascular endothelium) and TNC (expressed in stroma 2 and distal airway epithelium). *Robo1* knock-out mice exhibit delayed lung maturation with increased mesenchymal cellularity and reduced terminal air spaces leading to respiratory distress and death at birth^33^. TNC inactivation in mice induces abnormal lung development and persistent abnormal lung function after birth^34^. Lastly, we described crosstalk between immune cells and stroma populations via CD44-HAS2. CD44 plays an important role in the regulation of lung fibroblasts senescence and apoptosis and the interaction HAS2-CD44 can induce pulmonary fibrosis via its receptor ^35^. Immune cells population in our dataset were not well differentiated but additional data suggest that lung immune cells complexity and heterogeneity increase later during the gestation and after birth and play an important role in stroma remodelling and angiogenesis in the developing lung^36^.

Distal airway epithelial cells were among the most active signal-receiving cell types in our dataset, displaying active communication with all other cell populations including themselves. Our dataset further identifies a cluster of rare PNEC, and its interaction with both stroma 1 and 2, pericytes and vascular endothelium populations. The exact role of PNEC in lung development is yet to be described^37^. However, GRP expression had previously been detected in human fetal lung at 8 WG with peak of expression at mid-gestation. Moreover, elevated urine GRP was previously associated with the development of BPD suggesting an important role for PNECs in normal and impaired lung development ^38^. In addition, FISH identified 2 populations of PNEC within the fetal lung tissue (*GRP*^+^/*NRXN1*^+^and *GRP*^-^/*NRXN1*^+^ cells). These findings suggest that *NRXN1* might be a more specific marker for PNEC than *GRP*.

Our study has several limitations. Firstly, the study doesn’t consider the potential effects of fetus sex on the lung development whereas sex-related differences in human fetal lung transcriptome during the pseudoglandular stage had been previously reported^39^. However, the same study also found GA to have a more dominant effect on transcriptome than sex. Secondly, we assumed the fetuses were healthy as they were issued from selective abortions with no fetal indications. Nevertheless, all the fetuses had a GA below 20 weeks of gestation, prior to typical ultrasound morphological assessment for congenital malformation. Additional limitations relate to the downstream analysis. During development, important changes in cell differentiation and signalling can be induced by upregulation or downregulation in gene expression. While the presented DSA approach provides information on up-regulated genes across the different GA, it does not allow to study down-regulation in gene expression.

We report here, for the first time, an unbiased transcriptomic analysis of human fetal lungs during pseudoglandular stage at a single-cell resolution. This transcriptomic approach at a single cell level, combined with other novel approaches such as proteomics and metabolomics analysis^40,41^, are critical to unravel molecular pathways and cell communication in the human developing lung. Altogether, they will provide a clinically relevant background for the generation of novel research hypotheses in studies of normal or impaired lung development, help to develop and validate surrogate models to study human lung development and ultimately dentify new therapeutic targets.

