## supplemental method for "A single-cell atlas of human fetal lung development between 14 and 19 weeks of gestation"

**Supplemental Methods**

1. **Human fetal lung collection.**

De-identified human fetal lung samples were obtained from women undergoing elective termination of pregnancy with accurate dating (either last menstrual period or an ultrasound) and live, singleton pregnancy. Pregnancies with fetal morphological or chromosomal anomalies were excluded. Tissue collection was approved by The Ottawa Hospital Review Ethical Board (approval number 20170603-01H).

Lung samples were collected for precision-cut lung slices^1^. Briefly fetal lungs were sampled by the pathologist. Lungs were subsequently inflated with a low gelling temperature agarose solution (1.5% solution, #A9414 Sigma-Aldrich, Oakville, ON, Canada), and sampled with an 8mm biopsy punch (Robbins True-Cut Disposable Biopsy Punch 8mm, #RBP-80, Robbins instruments, Chatham, NJ, USA) prior to slicing with a tissue slicer. Spare lung tissue was snap frozen in liquid nitrogen and stored at -80°C for single nuclei isolation.

2. **Single nuclei isolation.**

Single nuclei isolation was performed according to Martelotto et al.^2^ with minor adjustments.

2.1 Tissue homogenisation.

From each fetal lung, 4 small pieces (each the size of a grain of rice) were randomly sampled. These 4 pieces were divided in two 1.5 ml tubes. The two pieces per 1.5 ml tube were mechanically dissociated on ice using a single-use plastic pestle (Bel-Art^®^ Disposable Polypropylene Pestles and 1.5ml tubes, #66001-104, VWR, Mont-Royal, QC, Canada). Homogenates were resuspended in 500 µl of chilled Nuclei EZ Lysis buffer (Nuclei Isolation Kit: Nuclei EZ Prep, #NUC101-1KT, Sigma-Aldrich, Oakville, ON, Canada). Samples corresponding to same fetal lung sample were pooled together and 500 µl of the homogenate was transferred to a new 2 ml tube.

2.2 Nuclei isolation, multiplexing and staining.

One milliliter of Nuclei EZ Lysis buffer (Nuclei Isolation Kit: Nuclei EZ Prep, #NUC101-1KT, Sigma-Aldrich, Oakville, ON, Canada) was added to each sample, mixed gently, and incubated on ice for 5 minutes. Homogenates were filtered through a 50µl nylon mesh (Thermofisher Scientific, Burlington, ON, Canada). Flow-through fractions were centrifuged at 500x g for 5 minutes.

Multiplexing was performed according to the MULTI-seq protocol^3^. The resulting pellets were re-suspended in 200 µl of 200 nM anchor/200 nM barcode solution, each sample receiving a different sample barcode. Samples were incubated on ice for 10 minutes, after which 16 µl of common lipid-modified co-anchor mix (2 µM) was added. Samples were incubated on ice for 5 minutes and diluted in 500 µl of Nuclei wash buffer (1x DPBS, Lonza, Basel, Switzerland) with 1% (v/w) Bovine Serum Albumin (Sigma-Aldrich, Oakville, ON, Canada) and 0.2 IU/µl RNase inhibitor (NProtector RNase inhibitor, #3335402001, Sigma-Aldrich, Oakville, ON, Canada). Single nuclei were counted with an automated cell counter (Eve Automatic cell Counter, Nano Entek, Waltham, MA, USA) and pelleted at 500× g for 5 minutes. Pellets were re-suspended with Nuclei Wash Buffer in order to achieve equal concentrations. Nuclei samples were pooled at equal ratio into a 1.5ml tube and pelleted at 500x g for 5 min. The pooled nuclei were washed with 500 µl of Nuclei wash buffer and pelleted at 500 x g for 5 minutes.

The pooled nuclei were re-suspended in 500 µl Nuclei Wash Buffer with DAPI (10 µg/ml). Nuclei integrity was confirmed using a fluorescent microscope (Axio Imager M2, Carl Zeiss, Toronto, ON, Canada).

2.3 Nuclei sorting.

Nuclei were sorted using a flow cytometer (BD LSR Fortessa, Beckton Dickinson Biosciences, Franklin Lakes, NJ, USA) at the Ottawa Hospital Research Institute (OHRI) core facility. Sample compensation was performed using Summit v.5.4 software and data analysis were performed with MoFlo XDP software (XDP, Beckman Coulter, Fullerton, CA, USA). The flow cytometer was gated on the single diploid DAPI peak, and 50,000 - 60,000 nuclei were sorted in a small volume of Nuclei Wash Buffer (~50 µl). Sorted nuclei were immediately processed by 10x Chromium at the OHRI StemCore facility.

3. **Single nuclei RNA sequencing and pre-processing.**

3.1 Library preparation and sequencing.

The 10x Chromium gene expression libraries were prepared according to manufacturer’s protocol (10X Genomics, Pleasanton, CA, USA), and the sequencing was performed at the OHRI StemCore facility using NextSeq500 (Illumina, San Diego, CA, USA).

3.2 Pre-processing.

Raw sequencing reads were processed using CellRanger v3.1.0, aligning reads to the mm10 build of the human genome. MULTI-seq barcode libraries were trimmed prior to demultiplexing using Trimmomatic (v0.36). Demultiplexing was performed using the deMULTIplex R package (v1.0.2)^3,4^. Only cells positive for a single barcode were kept for further analysis. Sample annotations were added to all cells in the data set. Pre-processing steps and downstream analysis were performed with Seurat (Seurat v4.0.0)^5^.

3.3 Quality control, normalization, and integration

Expression matrices were loaded as Seurat objects into R^6^. Only cells with > 200 genes detected and < 20% of UMIs mapped to mitochondrial genes were used. Each unique sample was split based on MULTI-seq sample barcodes into a separate Seurat object. SCTransform() was used to normalize samples, select highly variable genes, and to regress out cell cycle and cell stress effects. To eliminate batch effects or biological variability effects on clustering, the data integration method implemented by Seurat for SCTransform-normalized data was performed, using the SelectIntegrationFeatures(), PrepSCTIntegration(), FindIntegrationAnchors(), and IntegrateData() functions.

4. **Downstream analysis.**

4.1 Clustering and clusters annotation.

Dimensionality reduction was performed using principal component analysis (PCA) with RunPCA() function on the top 3000 variable genes and the data was clustered at a low resolution (dims: 20, resolution: 0.05, re-embedding dims: 20) with the Louvain algorithm implemented in the FindClusters() function in Seurat. Cell populations were identified with a simple Wilcoxon rank sum test with the FindAllMarkers() function in Seurat. For cluster annotation, we used a differential expression testing approach using FindAllMarkers() function to identify clusters marker genes (e.g., genes up-regulated in the cluster of interest). These marker genes were compared to known marker genes sets from LunGENS (Lung Gene Expression iN Single-cell) datasets on the LGEA (Lung Gene Expression Analysis) web portal^7,8^, and literature to define clusters identities. We next isolated and reprocessed the stromal and epithelial subsets using the same normalization and integration approach. In this case data were re-clustered at a higher resolution (Stroma dims: 20, resolution: 0.075, re-embedding dims:20; and for Distal airway epithelium dims: 20, resolution: 0.10, re-embedding dims:20). Upon subclustering the distal airway epithelium, we identified a small fourth subclustering showing high expression of stromal canonical markers. This population of cells was manually removed from the dataset. Gene set enrichment analysis (GSEA) was performed with Metascape^9^.

4.2 Differential State Analysis.

To identify differentially expressed genes in response to gestational age (GA), we performed a Differential State Analysis (DSA) using muscat R package^10^. Genes with an adjusted p-value <0.01 were considered significant and used to generate a heat map. From this heat map, genes were classified according to their up-regulated expression among different GA (individual samples).

To identify gene sets associated with differentially expressed genes we performed a GSEA using Metascape^9^. For each cluster, enrichment analysis was done using a single list (DSA results gene adj-p<0.01). In addition, we performed a meta-analysis using multiple gene list where each column represented a different GA and contained the up-regulated genes for this specific GA.

Differential gene expression across GA within a subpopulation was also used to identify potential early, late, and general (expressed across all the GA) marker genes for this subpopulation and further assess for spatial localization by FISH.

4.3 Cell communication

To describe and understand cell communication networks within the developing human fetal lung samples, we used NicheNet (v1.0.0) R package^11^. To prioritize results, analysis was limited to larger cell populations with higher number of differentially expressed genes across GA. Clusters 0, 1, 2, 3, 4, 5, and 6 were considered as signal receivers, while all the clusters were considered as potential signal senders. All the cell types were considered as sending cells, whereas ciliated cells and PNEC were not considered as receiving cell types as they represented small clusters with lack of changes in gene expression across the GA. In addition to crosstalk between diverse populations, crosstalk within the population itself was also observed in all the cell types, except PNEC and ciliated cells. To define a gene set of interest, the two latest GA timepoints (18+2w and 19+0w) were used as a condition of interest and two earliest GA timepoints (14+1w and 14+3w) as the reference condition. Background expression of genes was specified using all genes with adjusted p-value < 0.05, log2FC value > 0.05, and >10% detection in a given cluster. For each “receiver” cell population, top 20 ligands predicted to drive developmental age were selected based on the Pearson correlation coefficient. Quantile cut-off on the ligand-target scores of the input weighted ligand-target network was set to 0.33. To further validate the clinical relevance of our cell communication results, ligands expression in sending cell types and receptors expression in receiving cell types were compared to the Human Protein Atlas ([https://www.proteinatlas.org](https://www.proteinatlas.org/)) to confirm the expression of the protein (ligand or receptor) within the cell type of interest in human lung. Cell communication results are presented as a circos plot in Figure 6, and bona-fide receptors and predicted target genes for the receiving cell types are presented in Figure 7. When the predicted target genes list contained 3 or more genes, a GSEA was performed, and the results are presented in supplemental table 9.

5. **Data and code availability.**

All RNA sequencing data including raw fastq sequencing files, gene expression matrices, and cell metadata reported in this article is NCBI Gene Expression Omnibus (GEO) database and can be made available by request to the corresponding author. Code used for the analysis of the snRNA-seq data will be made available at the public GitHub repository and is available by request to the corresponding author.

6. **Fluorescent in situ hybridization.**

RNA in situ hybridization was performed on fresh 4 μm formalin-fixed paraffin embedded (FFPE) tissue sections using RNAscope Multiplex Fluorescent Reagent Kit v2 (#323100, Advanced Cell Diagnostics, CA, USA) according to the manufacturers’s instructions. Briefly, tissue sections were baked for 1 h at 60°C, deparaffinized and treated with hydrogen peroxide for 10 min at room temperature (RT). Target retrieval was performed for 15 min at 98°C, followed by protease plus treatment for 15 min at 40°C. The sections were then hybridized with probes for 2 h at 40°C followed by signal amplification and developing of HRP channels. The RNAscope probes used in this study were: Hs CAV1 (#452071), Hs FGFR2 (#311171-C2), Hs GRP (#465261) with 1:5 dilution, Hs KITLG (#407671-C3), Hs NRXN1 (527151-C3), Hs PDGFRB (#548991), Hs PECAM1-O1 (#455931-C2), Hs SFTBP-O1 (#1087181-C3), Hs SEMA3C (#549241), 3-plex positive control probe Hs (#320861), and 3-plex negative control probe (#320871). The signals were detected with TSA Plus fluorophores fluorescein (1:750 dilution), Cyanine 3 (1:1500 dilution), and Cyanine 5 (1:3000 dilution) (NEL741001KT, NEL744001KT, and NEL745001KT, respectively, Akoya Biosciences, MA, USA). The sections were counterstained with DAPI and mounted with ProLong Gold Antifade Mountant (P36930, Life Technologies Limited, Thermo Fisher Scientific, UK). Tissue sections were scanned using 3DHISTECH Pannoramic 250 FLASH II digital slide scanner at 40x magnification with extended focus and 7 focus levels at Genome Biology Unit supported by HiLIFE and the Faculty of Medicine, University of Helsinki, and Biocenter Finland.

**References.**

1	Kang MH, van Lieshout LP, Xu L, Domm JM, Vadivel A, Renesme L *et al.* A lung tropic AAV vector improves survival in a mouse model of surfactant B deficiency. *Nat Commun* 2020; **11**: 3929.

2	Martelotto L. ‘Frankenstein’ protocol for nuclei isolation from fresh and frozen tissue for snRNAseq. 2020. doi:10.17504/protocols.io.3fkgjkw.

3	McGinnis CS, Patterson DM, Winkler J, Conrad DN, Hein MY, Srivastava V *et al.* MULTI-seq: sample multiplexing for single-cell RNA sequencing using lipid-tagged indices. *Nat Methods* 2019; **16**: 619–626.

4	Hurskainen M, Mižíková I, Cook DP, Andersson N, Cyr-Depauw C, Lesage F *et al.* Single cell transcriptomic analysis of murine lung development on hyperoxia-induced damage. *Nat Commun* 2021; **12**: 1565.

5	Hao Y, Hao S, Andersen-Nissen E, Mauck WM, Zheng S, Butler A *et al.* Integrated analysis of multimodal single-cell data. *Cell* 2021; : S0092867421005833.

6	R Core Team (2020). R: A language and environment for statistical computing. R Foundation for Statistical Computing, Vienna, Austria. https://www.r-project.org/ (accessed 10 May2021).

7	Du Y, Guo M, Whitsett JA, Xu Y. ‘LungGENS’: a web-based tool for mapping single-cell gene expression in the developing lung. *Thorax* 2015; **70**: 1092–1094.

8	Du Y, Kitzmiller JA, Sridharan A, Perl AK, Bridges JP, Misra RS *et al.* Lung Gene Expression Analysis (LGEA): an integrative web portal for comprehensive gene expression data analysis in lung development. *Thorax* 2017; **72**: 481–484.

9	Zhou Y, Zhou B, Pache L, Chang M, Khodabakhshi AH, Tanaseichuk O *et al.* Metascape provides a biologist-oriented resource for the analysis of systems-level datasets. *Nat Commun* 2019; **10**: 1523.

10	Crowell HL, Soneson C, Germain P-L, Calini D, Collin L, Raposo C *et al.* muscat detects subpopulation-specific state transitions from multi-sample multi-condition single-cell transcriptomics data. *Nat Commun* 2020; **11**: 6077.

11	Browaeys R, Saelens W, Saeys Y. NicheNet: modeling intercellular communication by linking ligands to target genes. *Nat Methods* 2020; **17**: 159–162.
