## supplemental figures for "A single-cell atlas of human fetal lung development between 14 and 19 weeks of gestation"

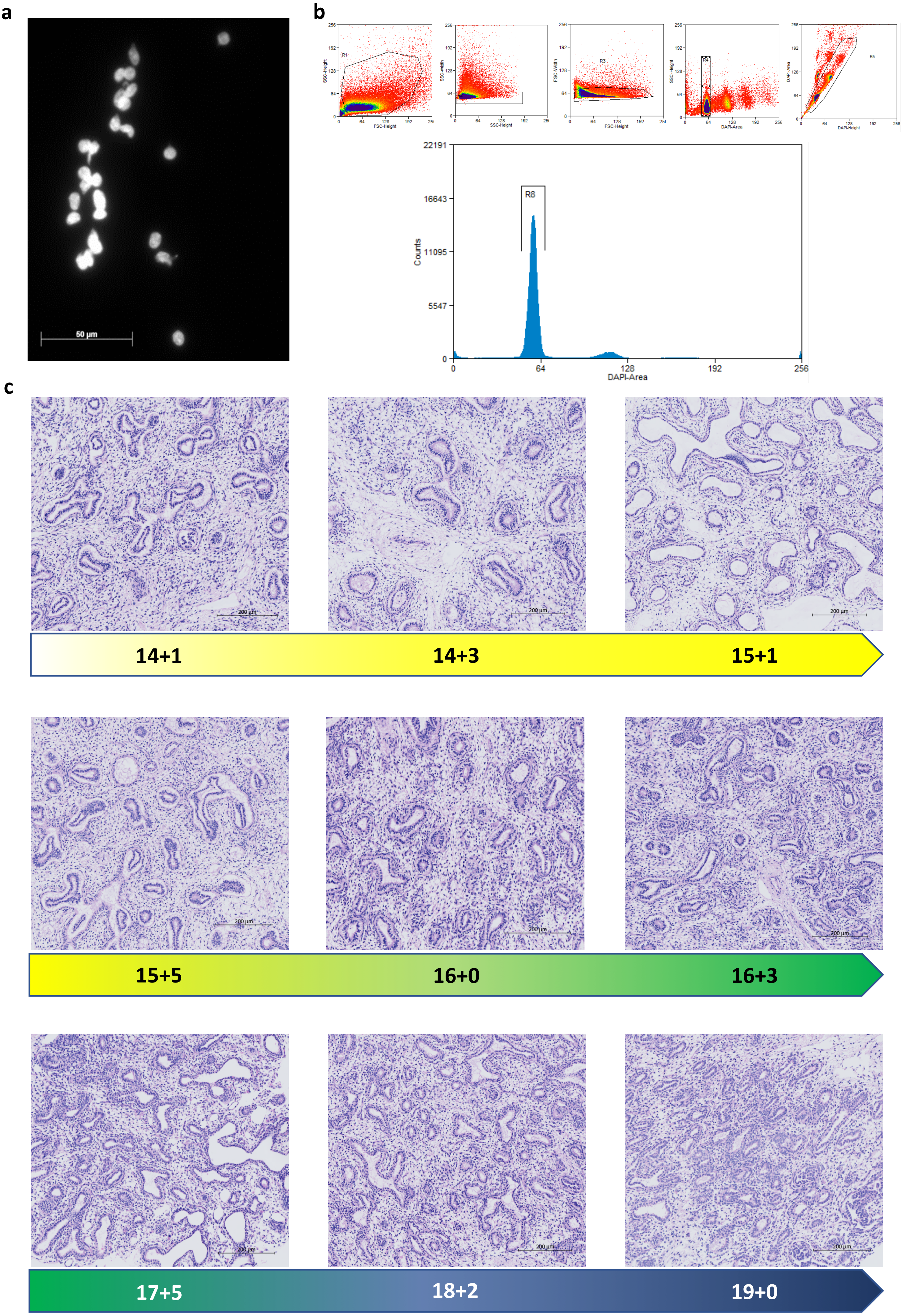
a
50 µm
b
R8
Counts
22191
16643
11095
5547
0
64
128
192
256
DAPI-Area
c
200 µm
14+1
14+3
15+1
15+5
16+0
16+3
17+5
18+2
19+0

**Supplemental figure 1. Quality control of the nuclei used for snRNA-seq and morphology of the human fetal lung samples**

a) Representative image showing isolated single nuclei stained with DAPI and visualized by fluorescent microscopy. Magnification x40. b) Gating strategy to obtain single diploid nuclei and histogram showing the purity and quality of the single nuclei suspension as evaluated by flow cytometry. c) Representative images showing the lung architecture in all fetal samples stained with hematoxylin and eosin. Magnification x20, scale = 200 µm.

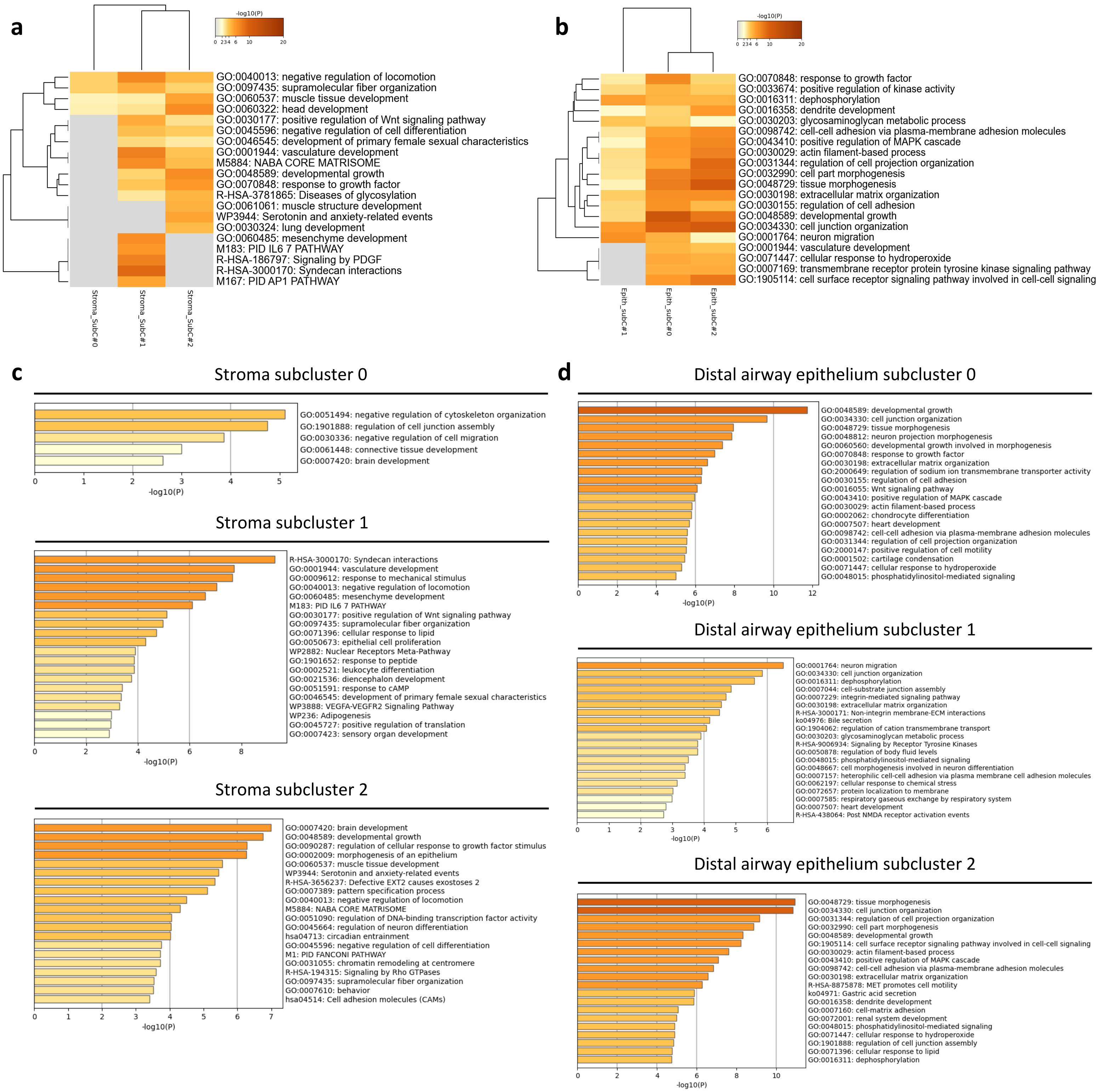
a
-log10(P)
0 2 3 4 6 10 20
GO:0040013: negative regulation of locomotion
GO:0097435: supramolecular fiber organization
GO:0060537: muscle tissue development
GO:0060322: head development
GO:0030177: positive regulation of Wnt signaling pathway
GO:0045596: negative regulation of cell differentiation
GO:0046545: development of primary female sexual characteristics
GO:0001944: vasculature development
M5884: NABA CORE MATRISOME
GO:0048589: developmental growth
GO:0070848: response to growth factor
R-HSA-3781865: Diseases of glycosylation
GO:0061061: muscle structure development
WP3944: Serotonin and anxiety-related events
GO:0030324: lung development
GO:0060485: mesenchyme development
M183: PID IL6 7 PATHWAY
R-HSA-186797: Signaling by PDGF
R-HSA-3000170: Syndecan interactions
M167: PID AP1 PATHWAY
Stroma_SubC#0
Stroma_SubC#1
Stroma_SubC#2
b
-log10(P)
0 2 3 4 6 10 20
GO:0070848: response to growth factor
GO:0033674: positive regulation of kinase activity
GO:0016311: dephosphorylation
GO:0016358: dendrite development
GO:0030203: glycosaminoglycan metabolic process
GO:0098742: cell-cell adhesion via plasma-membrane adhesion molecules
GO:0043410: positive regulation of MAPK cascade
GO:0030029: actin filament-based process
GO:0031344: regulation of cell projection organization
GO:0032990: cell part morphogenesis
GO:0048729: tissue morphogenesis
GO:0030198: extracellular matrix organization
GO:0030155: regulation of cell adhesion
GO:0048589: developmental growth
GO:0034330: cell junction organization
GO:0001764: neuron migration
GO:0001944: vasculature development
GO:0071447: cellular response to hydroperoxide
GO:0007169: transmembrane receptor protein tyrosine kinase signaling pathway
GO:1905114: cell surface receptor signaling pathway involved in cell-cell signaling
Epith_subC#1
Epith_subC#0
Epith_subC#2
c
Stroma subcluster 0
GO:0051494: negative regulation of cytoskeleton organization
GO:1901888: regulation of cell junction assembly
GO:0030336: negative regulation of cell migration
GO:0061448: connective tissue development
GO:0007420: brain development
0 1 2 3 4 5
-log10(P)
Stroma subcluster 1
R-HSA-3000170: Syndecan interactions
GO:0001944: vasculature development
GO:0009612: response to mechanical stimulus
GO:0040013: negative regulation of locomotion
GO:0060485: mesenchyme development
M183: PID IL6 7 PATHWAY
GO:0030177: positive regulation of Wnt signaling pathway
GO:0097435: supramolecular fiber organization
GO:0071396: cellular response to lipid
GO:0050673: epithelial cell proliferation
WP2882: Nuclear Receptors Meta-Pathway
GO:1901652: response to peptide
GO:0002521: leukocyte differentiation
GO:0021536: diencephalon development
GO:0051591: response to cAMP
GO:0046545: development of primary female sexual characteristics
WP3888: VEGFA-VEGFR2 Signaling Pathway
WP236: Adipogenesis
GO:0045727: positive regulation of translation
GO:0007423: sensory organ development
0 2 4 6 8
-log10(P)
Stroma subcluster 2
GO:0007420: brain development
GO:0048589: developmental growth
GO:0090287: regulation of cellular response to growth factor stimulus
GO:0002009: morphogenesis of an epithelium
GO:0060537: muscle tissue development
WP3944: Serotonin and anxiety-related events
R-HSA-3656237: Defective EXT2 causes exostoses 2
GO:0007389: pattern specification process
GO:0040013: negative regulation of locomotion
M5884: NABA CORE MATRISOME
GO:0051090: regulation of DNA-binding transcription factor activity
GO:0045664: regulation of neuron differentiation
hsa04713: circadian entrainment
GO:0045596: negative regulation of cell differentiation
M1: PID FANCONI PATHWAY
GO:0031055: chromatin remodeling at centromere
R-HSA-194315: Signaling by Rho GTPases
GO:0097435: supramolecular fiber organization
GO:0007610: behavior
hsa04514: Cell adhesion molecules (CAMs)
0 1 2 3 4 5 6 7
-log10(P)
d
Distal airway epithelium subcluster 0
GO:0048589: developmental growth
GO:0034330: cell junction organization
GO:0048729: tissue morphogenesis
GO:0048812: neuron projection morphogenesis
GO:0060560: developmental growth involved in morphogenesis
GO:0070848: response to growth factor
GO:0030198: extracellular matrix organization
GO:2000649: regulation of sodium ion transmembrane transporter activity
GO:0030155: regulation of cell adhesion
GO:0016055: Wnt signaling pathway
GO:0043410: positive regulation of MAPK cascade
GO:0030029: actin filament-based process
GO:0002062: chondrocyte differentiation
GO:0007507: heart development
GO:0098742: cell-cell adhesion via plasma-membrane adhesion molecules
GO:0031344: regulation of cell projection organization
GO:2000147: positive regulation of cell motility
GO:0001502: cartilage condensation
GO:0071447: cellular response to hydroperoxide
GO:0048015: phosphatidylinositol-mediated signaling
0 2 4 6 8 10 12
-log10(P)
Distal airway epithelium subcluster 1
GO:0001764: neuron migration
GO:0034330: cell junction organization
GO:0016311: dephosphorylation
GO:0007044: cell-substrate junction assembly
GO:0007229: integrin-mediated signaling pathway
GO:0030198: extracellular matrix organization
R-HSA-3000171: Non-integrin membrane-ECM interactions
ko04976: Bile secretion
GO:1904062: regulation of cation transmembrane transport
GO:0030203: glycosaminoglycan metabolic process
R-HSA-9006934: Signaling by Receptor Tyrosine Kinases
GO:0050878: regulation of body fluid levels
GO:0048015: phosphatidylinositol-mediated signaling
GO:0048667: cell morphogenesis involved in neuron differentiation
GO:0007157: heterophilic cell-cell adhesion via plasma membrane cell adhesion molecules
GO:0062197: cellular response to chemical stress
GO:0072657: protein localization to membrane
GO:0007585: respiratory gaseous exchange by respiratory system
GO:0007507: heart development
R-HSA-438064: Post NMDA receptor activation events
0 1 2 3 4 5 6
-log10(P)
Distal airway epithelium subcluster 2
GO:0048729: tissue morphogenesis
GO:0034330: cell junction organization
GO:0031344: regulation of cell projection organization
GO:0032990: cell part morphogenesis
GO:0048589: developmental growth
GO:1905114: cell surface receptor signaling pathway involved in cell-cell signaling
GO:0030029: actin filament-based process
GO:0043410: positive regulation of MAPK cascade
GO:0098742: cell-cell adhesion via plasma-membrane adhesion molecules
GO:0030198: extracellular matrix organization
R-HSA-8875878: MET promotes cell motility
ko04971: Gastric acid secretion
GO:0016358: dendrite development
GO:0007160: cell-matrix adhesion
GO:0072001: renal system development
GO:0048015: phosphatidylinositol-mediated signaling
GO:0071447: cellular response to hydroperoxide
GO:1901888: regulation of cell junction assembly
GO:0071396: cellular response to lipid
GO:0016311: dephosphorylation
0 2 4 6 8 10
-log10(P)

**Supplemental figure 2. Gene set enrichment for stromal and distal airway epithelial subclusters**

a) Heatmaps depicting the enriched terms associated with different stromal subclusters as identified by multi-list enrichment analysis. b) Heatmaps depicting the enriched terms associated with different distal airway epithelium subclusters as identified by multi-list enrichment analysis. c) Enriched terms associated with individual stromal subclusters as identified by enrichment analysis. d) Enriched terms associated with individual distal airway epithelium subclusters as identified by enrichment analysis.

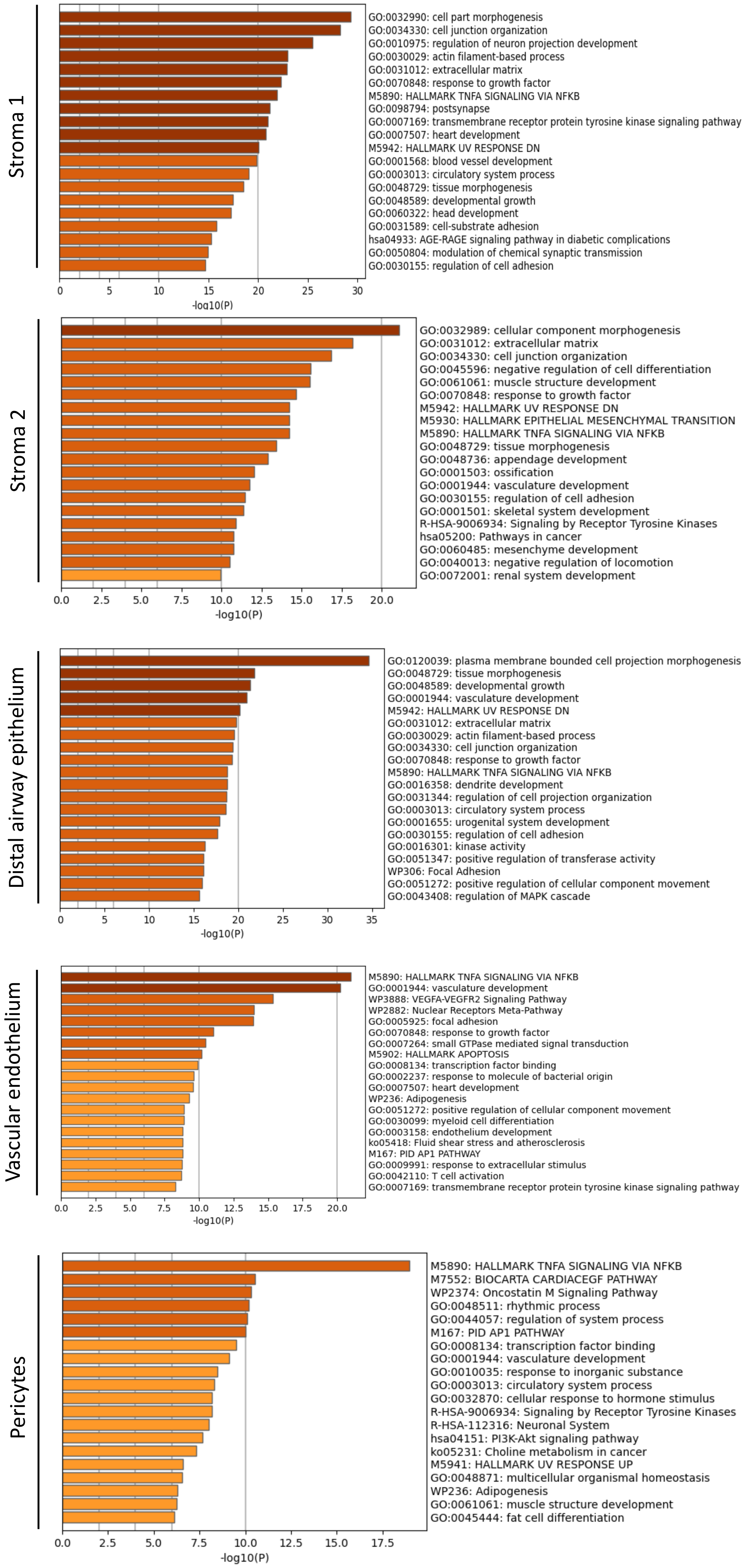
Stroma 1
GO:0032990: cell part morphogenesis
GO:0034330: cell junction organization
GO:0010975: regulation of neuron projection development
GO:0030029: actin filament-based process
GO:0031012: extracellular matrix
GO:0070848: response to growth factor
M5890: HALLMARK TNFA SIGNALING VIA NFKB
GO:0098794: postsynapse
GO:0007169: transmembrane receptor protein tyrosine kinase signaling pathway
GO:0007507: heart development
M5942: HALLMARK UV RESPONSE DN
GO:0001568: blood vessel development
GO:0003013: circulatory system process
GO:0048729: tissue morphogenesis
GO:0048589: developmental growth
GO:0060322: head development
GO:0031589: cell-substrate adhesion
hsa04933: AGE-RAGE signaling pathway in diabetic complications
GO:0050804: modulation of chemical synaptic transmission
GO:0030155: regulation of cell adhesion
0 5 10 15 20 25 30
-log10(P)
Stroma 2
GO:0032989: cellular component morphogenesis
GO:0031012: extracellular matrix
GO:0034330: cell junction organization
GO:0045596: negative regulation of cell differentiation
GO:0061061: muscle structure development
GO:0070848: response to growth factor
M5942: HALLMARK UV RESPONSE DN
M5930: HALLMARK EPITHELIAL MESENCHYMAL TRANSITION
M5890: HALLMARK TNFA SIGNALING VIA NFKB
GO:0048729: tissue morphogenesis
GO:0048736: appendage development
GO:0001503: ossification
GO:0001944: vasculature development
GO:0030155: regulation of cell adhesion
GO:0001501: skeletal system development
R-HSA-9006934: Signaling by Receptor Tyrosine Kinases
hsa05200: Pathways in cancer
GO:0060485: mesenchyme development
GO:0040013: negative regulation of locomotion
GO:0072001: renal system development
0.0 2.5 5.0 7.5 10.0 12.5 15.0 17.5 20.0
-log10(P)
Distal airway epithelium
GO:0120039: plasma membrane bounded cell projection morphogenesis
GO:0048729: tissue morphogenesis
GO:0048589: developmental growth
GO:0001944: vasculature development
M5942: HALLMARK UV RESPONSE DN
GO:0031012: extracellular matrix
GO:0030029: actin filament-based process
GO:0034330: cell junction organization
GO:0070848: response to growth factor
M5890: HALLMARK TNFA SIGNALING VIA NFKB
GO:0016358: dendrite development
GO:0031344: regulation of cell projection organization
GO:0003013: circulatory system process
GO:0001655: urogenital system development
GO:0030155: regulation of cell adhesion
GO:0016301: kinase activity
GO:0051347: positive regulation of transferase activity
WP306: Focal Adhesion
GO:0051272: positive regulation of cellular component movement
GO:0043408: regulation of MAPK cascade
0 5 10 15 20 25 30 35
-log10(P)
Vascular endothelium
M5890: HALLMARK TNFA SIGNALING VIA NFKB
GO:0001944: vasculature development
WP3888: VEGFA-VEGFR2 Signaling Pathway
WP2882: Nuclear Receptors Meta-Pathway
GO:0005925: focal adhesion
GO:0070848: response to growth factor
GO:0007264: small GTPase mediated signal transduction
M5902: HALLMARK APOPTOSIS
GO:0008134: transcription factor binding
GO:0002237: response to molecule of bacterial origin
GO:0007507: heart development
WP236: Adipogenesis
GO:0051272: positive regulation of cellular component movement
GO:0030099: myeloid cell differentiation
GO:0003158: endothelium development
ko05418: Fluid shear stress and atherosclerosis
M167: PID AP1 PATHWAY
GO:0009991: response to extracellular stimulus
GO:0042110: T cell activation
GO:0007169: transmembrane receptor protein tyrosine kinase signaling pathway
0.0 2.5 5.0 7.5 10.0 12.5 15.0 17.5 20.0
-log10(P)
Pericytes
M5890: HALLMARK TNFA SIGNALING VIA NFKB
M7552: BIOCARTA CARDIACEGF PATHWAY
WP2374: Oncostatin M Signaling Pathway
GO:0048511: rhythmic process
GO:0044057: regulation of system process
M167: PID AP1 PATHWAY
GO:0008134: transcription factor binding
GO:0001944: vasculature development
GO:0010035: response to inorganic substance
GO:0003013: circulatory system process
GO:0032870: cellular response to hormone stimulus
R-HSA-9006934: Signaling by Receptor Tyrosine Kinases
R-HSA-112316: Neuronal System
hsa04151: PI3K-Akt signaling pathway
ko05231: Choline metabolism in cancer
M5941: HALLMARK UV RESPONSE UP
GO:0048871: multicellular organismal homeostasis
WP236: Adipogenesis
GO:0061061: muscle structure development
GO:0045444: fat cell differentiation
0.0 2.5 5.0 7.5 10.0 12.5 15.0 17.5
-log10(P)

**Supplemental figure 3. Pathways induced by increasing age between 14 and 19 weeks of gestation in the fetal lung stromal, distal airway epithelium, vascular endothelium and pericytes populations**
Pathways induced during fetal lung development (14+1 until 19+0 weeks) as identified by enrichment analysis based on the DSA analysis in stroma 1, stroma 2, distal airway epithelium, vascular endothelium and lung pericytes.

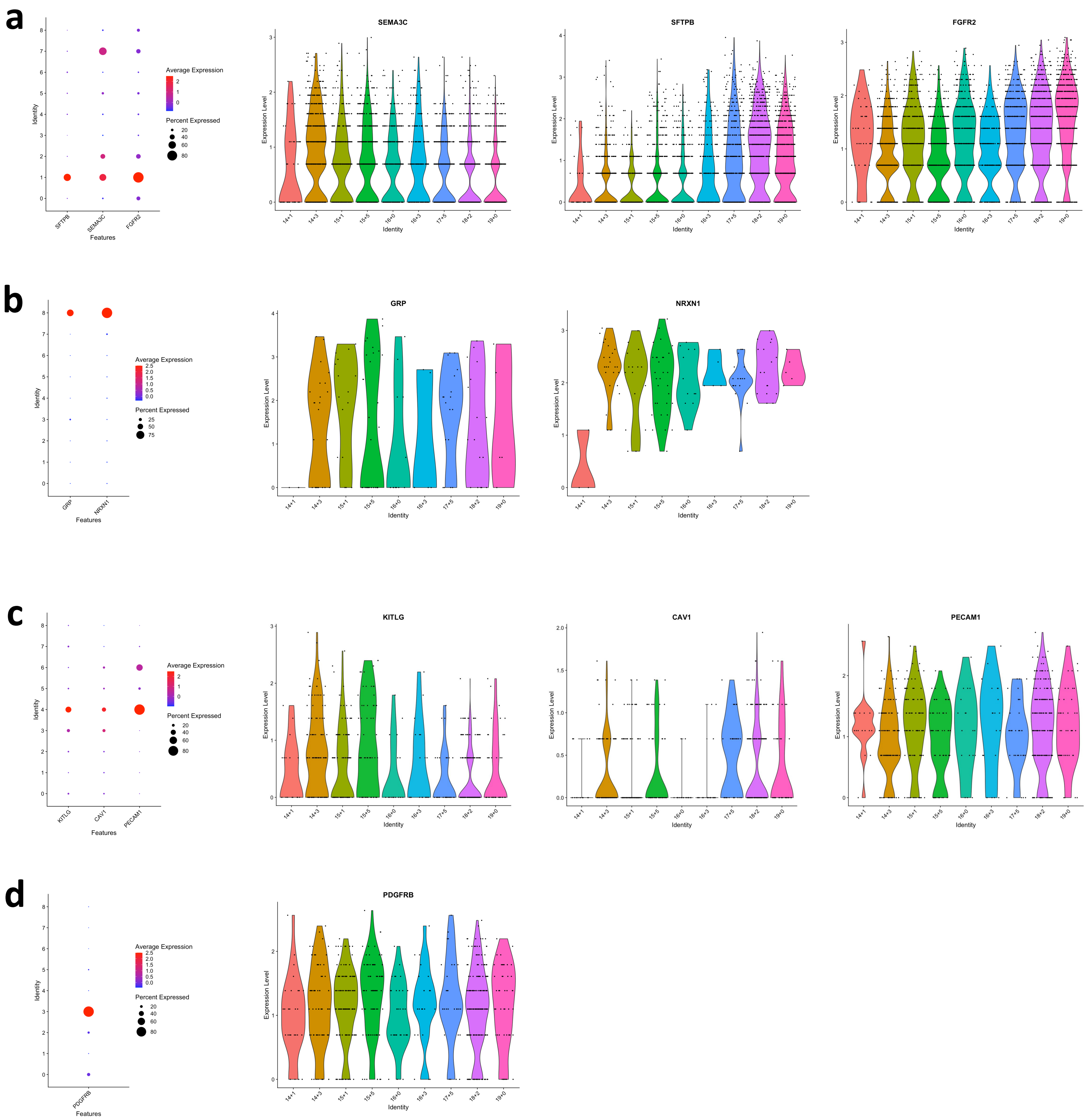
a
Identity
Features
SFTPB
SEMA3C
FGFR2
Average Expression
Percent Expressed
Expression Level
14+1
14+3
15+1
15+5
16+0
16+3
17+5
18+2
19+0
b
GRP
NRXN1
c
KITLG
CAV1
PECAM1
d
PDGFRB

**Supplemental figure 4. Gene selection for spatial validation by Fluorescent *In Situ* Hybridization (FISH)**

a) Dotplot depicting distal airway epithelium cluster specificity for *FGFR2*, *SEMA3C* and *SFTPB* (left). Violin plots (right) depicting gene expression level across different GAs (14+1 to 19+0 weeks of gestation, x-axis) and identifying *SEMA3C* as an early marker, *SFTPB* as a late marker and *FGFR2* as a general marker of the distal airway epithelium cluster. b) Dotplot depicting pulmonary neuroendocrine cell cluster specificity for *GRP* and *NRXN1* (left). Violin plot (right) depicting *GRP* and *NRXN1* expression level across different GAs (14+1 to 19+0 weeks of gestation, x-axis) and identifying *GRP* and *NRXN1* as a general marker of the pulmonary neuroendocrine cell cluster. Expression values in violin plots represent Z-score-transformed log(TP10k+1) values. c) Dotplot depicting vascular endothelium cluster specificity for *KITLG*, *CAV1* and *PECAM1* (left). Violin plots (right) depicting gene expression level across different GAs (14+1 to 19+0 weeks of gestation, x-axis) and identifying *KTLIG* as an early marker, *CAV1* as a late marker and *PECAM1* as a general marker of the vascular endothelium cluster. d) Dotplot depicting pericyte cluster specificity for *PDGFRB* (left). Violin plot (right) depicting *PDGFRB* expression level across different GAs (14+1 to 19+0 weeks of gestation, x-axis) and identifying *PDGFRB* as a general marker of the pericyte cluster.

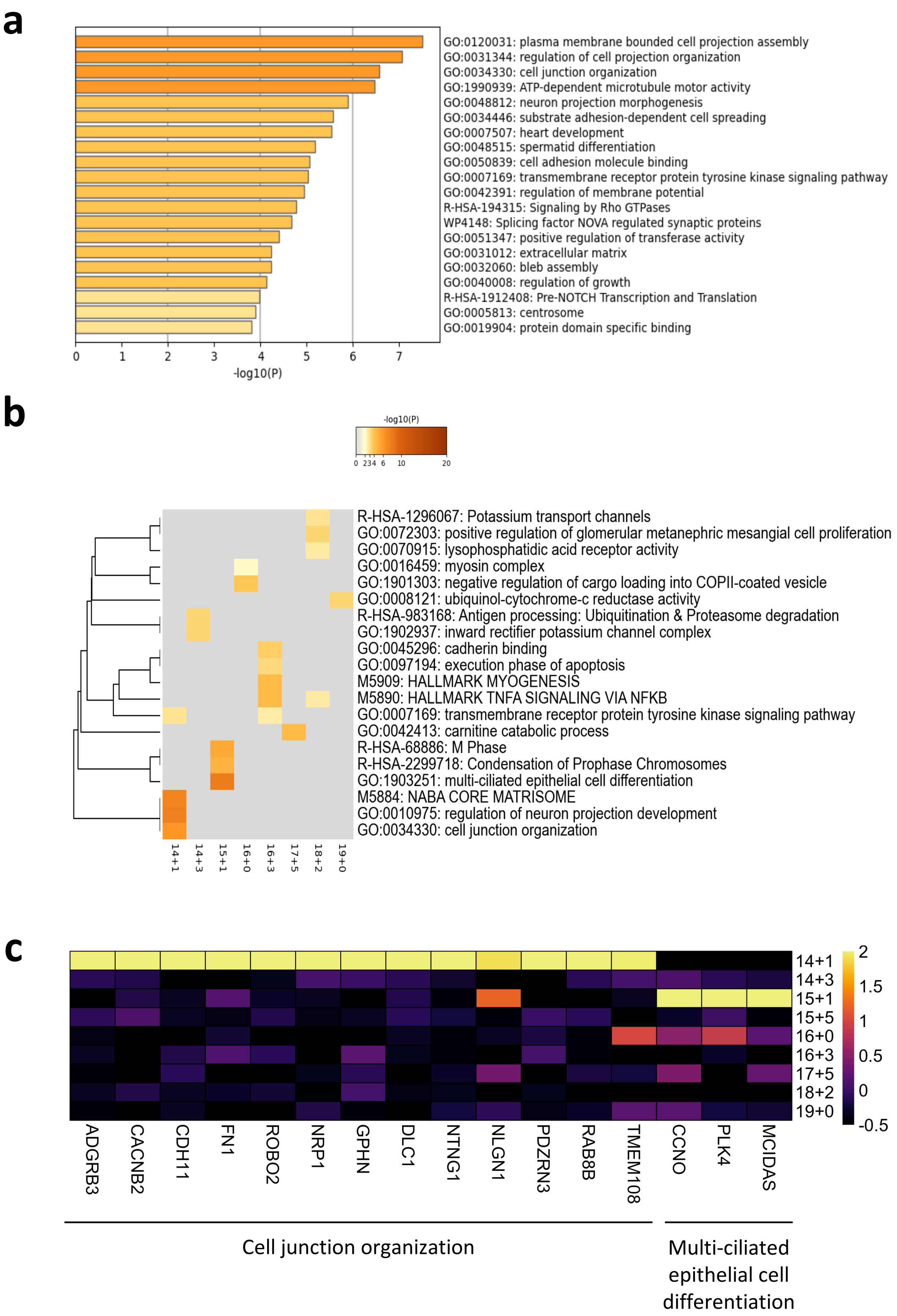
a
GO:0120031: plasma membrane bounded cell projection assembly
GO:0031344: regulation of cell projection organization
GO:0034330: cell junction organization
GO:1990939: ATP-dependent microtubule motor activity
GO:0048812: neuron projection morphogenesis
GO:0034446: substrate adhesion-dependent cell spreading
GO:0007507: heart development
GO:0048515: spermatid differentiation
GO:0050839: cell adhesion molecule binding
GO:0007169: transmembrane receptor protein tyrosine kinase signaling pathway
GO:0042391: regulation of membrane potential
R-HSA-194315: Signaling by Rho GTPases
WP4148: Splicing factor NOVA regulated synaptic proteins
GO:0051347: positive regulation of transferase activity
GO:0031012: extracellular matrix
GO:0032060: bleb assembly
GO:0040008: regulation of growth
R-HSA-1912408: Pre-NOTCH Transcription and Translation
GO:0005813: centrosome
GO:0019904: protein domain specific binding
0 1 2 3 4 5 6 7
-log10(P)
b
-log10(P)
0 234 6 10 20
R-HSA-1296067: Potassium transport channels
GO:0072303: positive regulation of glomerular metanephric mesangial cell proliferation
GO:0070915: lysophosphatidic acid receptor activity
GO:0016459: myosin complex
GO:1901303: negative regulation of cargo loading into COPII-coated vesicle
GO:0008121: ubiquinol-cytochrome-c reductase activity
R-HSA-983168: Antigen processing: Ubiquitination & Proteasome degradation
GO:1902937: inward rectifier potassium channel complex
GO:0045296: cadherin binding
GO:0097194: execution phase of apoptosis
M5909: HALLMARK MYOGENESIS
M5890: HALLMARK TNFA SIGNALING VIA NFKB
GO:0007169: transmembrane receptor protein tyrosine kinase signaling pathway
GO:0042413: carnitine catabolic process
R-HSA-68886: M Phase
R-HSA-2299718: Condensation of Prophase Chromosomes
GO:1903251: multi-ciliated epithelial cell differentiation
M5884: NABA CORE MATRISOME
GO:0010975: regulation of neuron projection development
GO:0034330: cell junction organization
14+1 14+3 15+1 16+0 16+3 17+5 18+2 19+0
c
14+1
14+3
15+1
15+5
16+0
16+3
17+5
18+2
19+0
2 1.5 1 0.5 0 -0.5
ADGRB3 CACNB2 CDH11 FN1 ROBO2 NRP1 GPHN DLC1 NTNG1 NLGN1 PDZRN3 RAB8B TMEM108 CCNO PLK4 MCIDAS
Cell junction organization
Multi-ciliated epithelial cell differentiation

**Supplemental figure 5. Developmental changes in gene expression in the fetal lung ciliated cells**
a) Pathways induced during fetal lung development (14+1 until 19+0 weeks) in the ciliated cells as identified by enrichment analysis based on the DSA. b) Heatmaps depicting the enriched terms associated with individual GAs as identified by multi-list enrichment analysis based on DSA in immune cells. c) Heatmap depicting the expression level of genes associated with selected enriched terms as identified in panel b). Expression levels in the heatmaps and dotplots are presented as log(TP10k+1) values. Log(TP10k+1) corresponds to log-transformed UMIs per 10k.

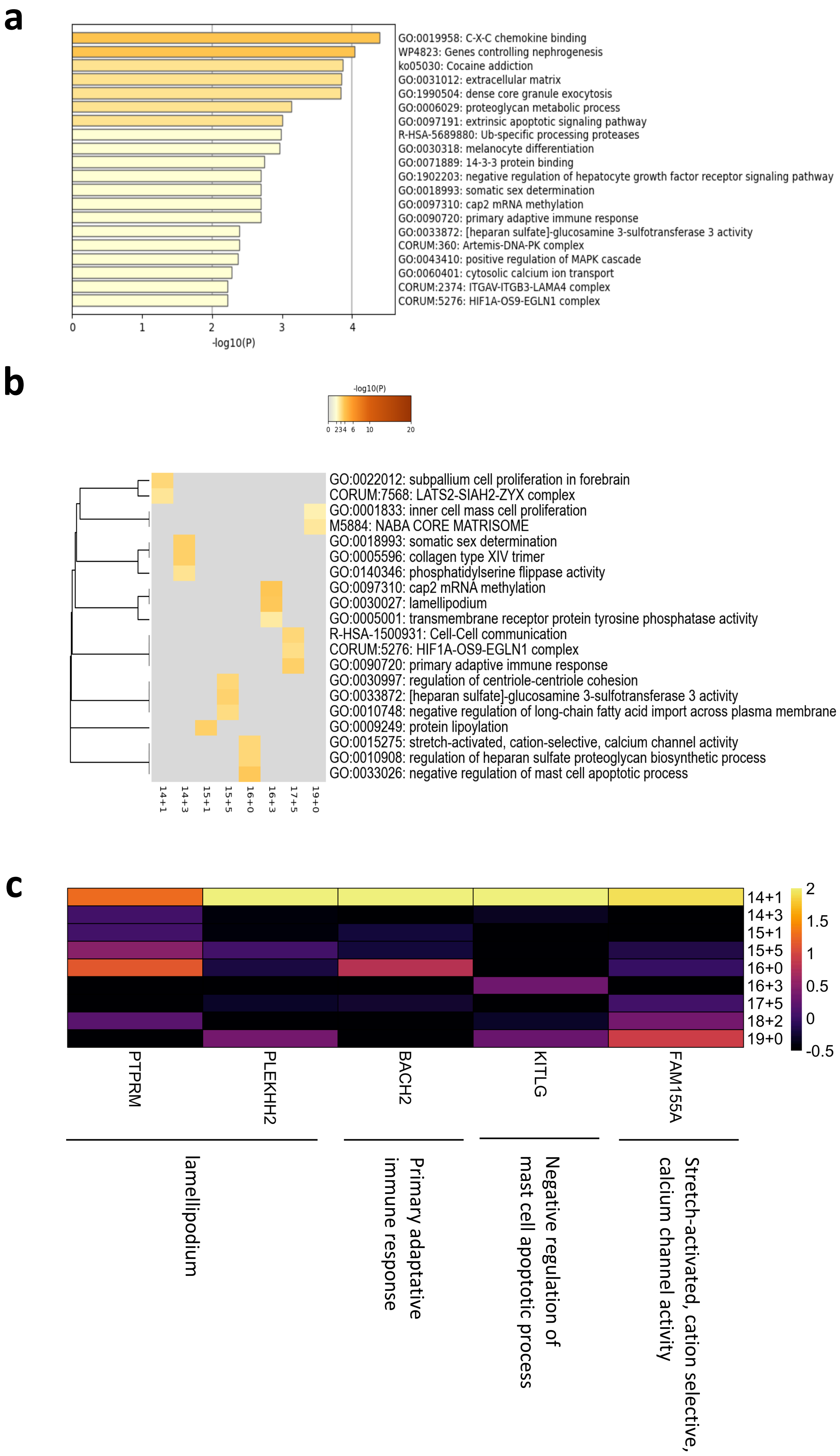
a
GO:0019958: C-X-C chemokine binding
WP4823: Genes controlling nephrogenesis
ko05030: Cocaine addiction
GO:0031012: extracellular matrix
GO:1990504: dense core granule exocytosis
GO:0006029: proteoglycan metabolic process
GO:0097191: extrinsic apoptotic signaling pathway
R-HSA-5689880: Ub-specific processing proteases
GO:0030318: melanocyte differentiation
GO:0071889: 14-3-3 protein binding
GO:1902203: negative regulation of hepatocyte growth factor receptor signaling pathway
GO:0018993: somatic sex determination
GO:0097310: cap2 mRNA methylation
GO:0090720: primary adaptive immune response
GO:0033872: [heparan sulfate]-glucosamine 3-sulfotransferase 3 activity
CORUM:360: Artemis-DNA-PK complex
GO:0043410: positive regulation of MAPK cascade
GO:0060401: cytosolic calcium ion transport
CORUM:2374: ITGAV-ITGB3-LAMA4 complex
CORUM:5276: HIF1A-OS9-EGLN1 complex
0
1
2
3
4
-log10(P)
b
-log10(P)
0 234 6 10 20
GO:0022012: subpallium cell proliferation in forebrain
CORUM:7568: LATS2-SIAH2-ZYX complex
GO:0001833: inner cell mass cell proliferation
M5884: NABA CORE MATRISOME
GO:0018993: somatic sex determination
GO:0005596: collagen type XIV trimer
GO:0140346: phosphatidylserine flippase activity
GO:0097310: cap2 mRNA methylation
GO:0030027: lamellipodium
GO:0005001: transmembrane receptor protein tyrosine phosphatase activity
R-HSA-1500931: Cell-Cell communication
CORUM:5276: HIF1A-OS9-EGLN1 complex
GO:0090720: primary adaptive immune response
GO:0030997: regulation of centriole-centriole cohesion
GO:0033872: [heparan sulfate]-glucosamine 3-sulfotransferase 3 activity
GO:0010748: negative regulation of long-chain fatty acid import across plasma membrane
GO:0009249: protein lipoylation
GO:0015275: stretch-activated, cation-selective, calcium channel activity
GO:0010908: regulation of heparan sulfate proteoglycan biosynthetic process
GO:0033026: negative regulation of mast cell apoptotic process
14+1
14+3
15+1
15+5
16+0
16+3
17+5
19+0
c
14+1
14+3
15+1
15+5
16+0
16+3
17+5
18+2
19+0
2
1.5
1
0.5
0
-0.5
PTPRM
PLEKHH2
BACH2
KITLG
FAM155A
lamellipodium
Primary adaptative immune response
Negative regulation of mast cell apoptotic process
Stretch-activated, cation selective, calcium channel activity

**Supplemental figure 6. Developmental changes in gene expression in the fetal lung pulmonary neuroendocrine cells**

a) Pathways induced during fetal lung development (14+1 until 19+0 weeks) in the pulmonary neuroendocrine cells (PNEC) as identified by enrichment analysis based on the DSA. b) Heatmaps depicting the enriched terms associated with individual GAs as identified by multi-list enrichment analysis based on DSA in immune cells. c) Heatmap depicting the expression level of genes associated with selected enriched terms as identified in panel b). Expression levels in the heatmaps and dotplots are presented as log(TP10k+1) values. Log(TP10k+1) corresponds to log-transformed UMIs per 10k.

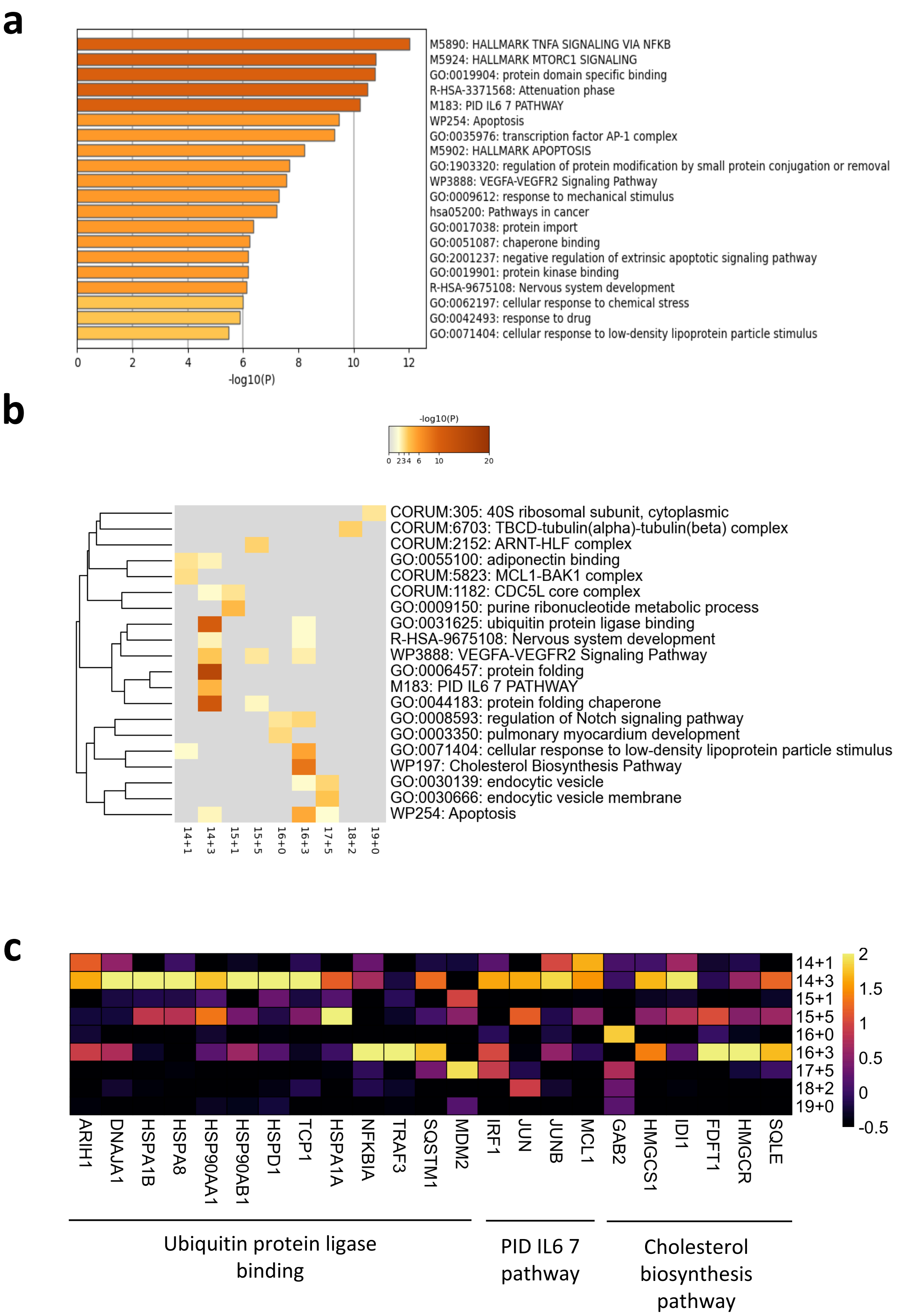

a
M5890: HALLMARK TNFA SIGNALING VIA NFKB
M5924: HALLMARK MTORC1 SIGNALING
GO:0019904: protein domain specific binding
R-HSA-3371568: Attenuation phase
M183: PID IL6 7 PATHWAY
WP254: Apoptosis
GO:0035976: transcription factor AP-1 complex
M5902: HALLMARK APOPTOSIS
GO:1903320: regulation of protein modification by small protein conjugation or removal
WP3888: VEGFA-VEGFR2 Signaling Pathway
GO:0009612: response to mechanical stimulus
hsa05200: Pathways in cancer
GO:0017038: protein import
GO:0051087: chaperone binding
GO:2001237: negative regulation of extrinsic apoptotic signaling pathway
GO:0019901: protein kinase binding
R-HSA-9675108: Nervous system development
GO:0062197: cellular response to chemical stress
GO:0042493: response to drug
GO:0071404: cellular response to low-density lipoprotein particle stimulus
0 2 4 6 8 10 12
-log10(P)
b
-log10(P)
0 2 3 4 6 10 20
CORUM:305: 40S ribosomal subunit, cytoplasmic
CORUM:6703: TBCD-tubulin(alpha)-tubulin(beta) complex
CORUM:2152: ARNT-HLF complex
GO:0055100: adiponectin binding
CORUM:5823: MCL1-BAK1 complex
CORUM:1182: CDC5L core complex
GO:0009150: purine ribonucleotide metabolic process
GO:0031625: ubiquitin protein ligase binding
R-HSA-9675108: Nervous system development
WP3888: VEGFA-VEGFR2 Signaling Pathway
GO:0006457: protein folding
M183: PID IL6 7 PATHWAY
GO:0044183: protein folding chaperone
GO:0008593: regulation of Notch signaling pathway
GO:0003350: pulmonary myocardium development
GO:0071404: cellular response to low-density lipoprotein particle stimulus
WP197: Cholesterol Biosynthesis Pathway
GO:0030139: endocytic vesicle
GO:0030666: endocytic vesicle membrane
WP254: Apoptosis
14+1 14+3 15+1 15+5 16+0 16+3 17+5 18+2 19+0
c
14+1
14+3
15+1
15+5
16+0
16+3
17+5
18+2
19+0
2 1.5 1 0.5 0 -0.5
ARIH1 DNAJA1 HSPA1B HSPA8 HSP90AA1 HSP90AB1 HSPD1 TCP1 HSPA1A NFKBIA TRAF3 SQSTM1 MDM2 IRF1 JUN JUNB MCL1 GAB2 HMGCS1 IDI1 FDFT1 HMGCR SQLE
Ubiquitin protein ligase binding
PID IL6 7 pathway
Cholesterol biosynthesis pathway

**Supplemental figure 7. Developmental changes in gene expression in the fetal lung lymphatic endothelial cells**

a) pathways induced during fetal lung development (14+1 until 19+0 weeks) in the lymphatic endothelial cells as identified by enrichment analysis based on the DSA. b) Heatmaps depicting the enriched terms associated with individual GAs as identified by multi-list enrichment analysis based on DSA in immune cells. c) Heatmap depicting the expression level of genes associated with selected enriched terms as identified in panel b). Expression levels in the heatmaps and dotplots are presented as log(TP10k+1) values. Log(TP10k+1) corresponds to log-transformed UMIs per 10k.

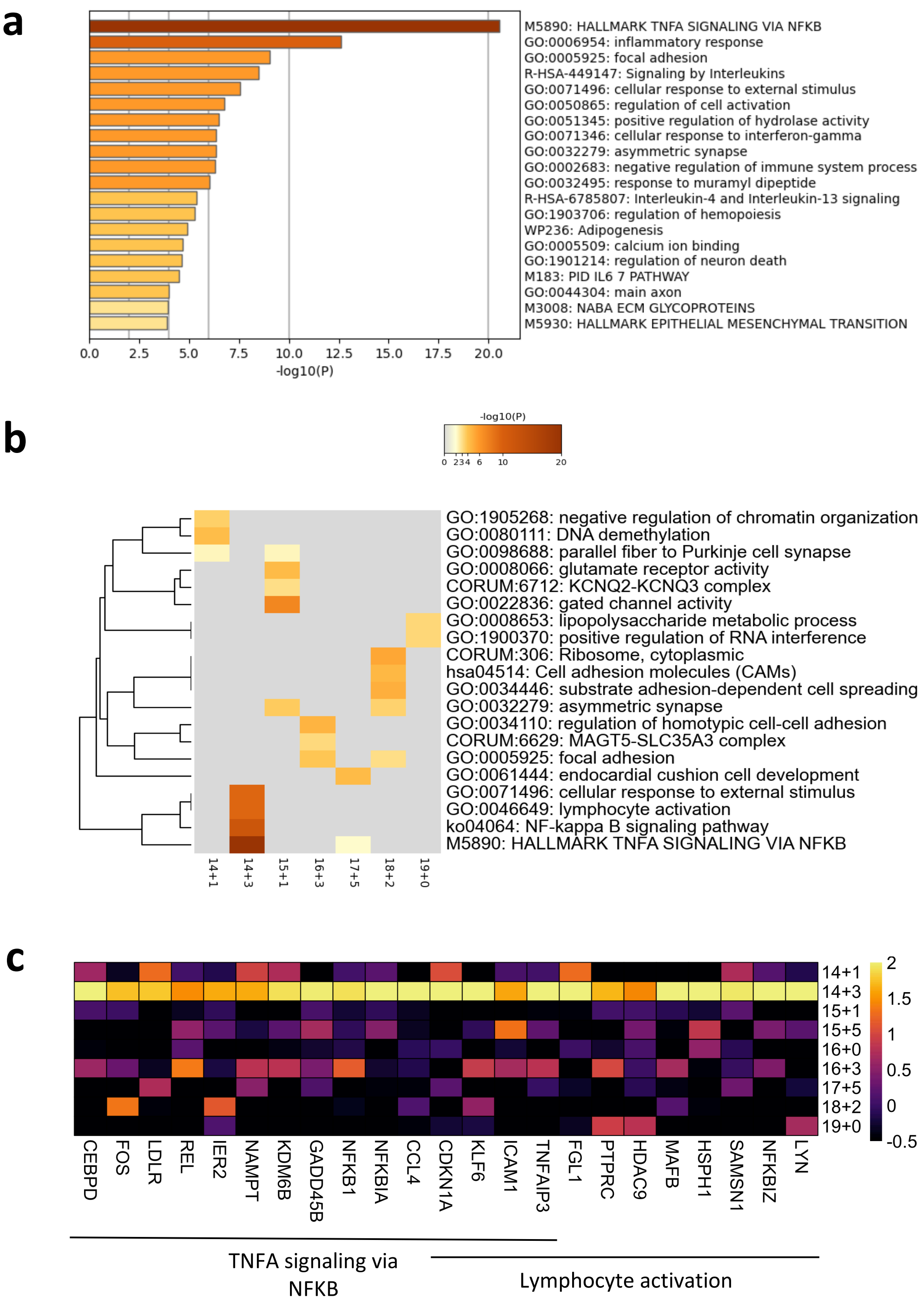

a
M5890: HALLMARK TNFA SIGNALING VIA NFKB
GO:0006954: inflammatory response
GO:0005925: focal adhesion
R-HSA-449147: Signaling by Interleukins
GO:0071496: cellular response to external stimulus
GO:0050865: regulation of cell activation
GO:0051345: positive regulation of hydrolase activity
GO:0071346: cellular response to interferon-gamma
GO:0032279: asymmetric synapse
GO:0002683: negative regulation of immune system process
GO:0032495: response to muramyl dipeptide
R-HSA-6785807: Interleukin-4 and Interleukin-13 signaling
GO:1903706: regulation of hemopoiesis
WP236: Adipogenesis
GO:0005509: calcium ion binding
GO:1901214: regulation of neuron death
M183: PID IL6 7 PATHWAY
GO:0044304: main axon
M3008: NABA ECM GLYCOPROTEINS
M5930: HALLMARK EPITHELIAL MESENCHYMAL TRANSITION
0.0 2.5 5.0 7.5 10.0 12.5 15.0 17.5 20.0
-log10(P)
b
-log10(P)
0 2 3 4 6 10 20
GO:1905268: negative regulation of chromatin organization
GO:0080111: DNA demethylation
GO:0098688: parallel fiber to Purkinje cell synapse
GO:0008066: glutamate receptor activity
CORUM:6712: KCNQ2-KCNQ3 complex
GO:0022836: gated channel activity
GO:0008653: lipopolysaccharide metabolic process
GO:1900370: positive regulation of RNA interference
CORUM:306: Ribosome, cytoplasmic
hsa04514: Cell adhesion molecules (CAMs)
GO:0034446: substrate adhesion-dependent cell spreading
GO:0032279: asymmetric synapse
GO:0034110: regulation of homotypic cell-cell adhesion
CORUM:6629: MAGT5-SLC35A3 complex
GO:0005925: focal adhesion
GO:0061444: endocardial cushion cell development
GO:0071496: cellular response to external stimulus
GO:0046649: lymphocyte activation
ko04064: NF-kappa B signaling pathway
M5890: HALLMARK TNFA SIGNALING VIA NFKB
14+1 14+3 15+1 16+3 17+5 18+2 19+0
c
14+1
14+3
15+1
15+5
16+0
16+3
17+5
18+2
19+0
2 1.5 1 0.5 0 -0.5
CEBPD FOS LDLR REL IER2 NAMPT KDM6B GADD45B NFKB1 NFKBIA CCL4 CDKN1A KLF6 ICAM1 TNFAIP3 FGL1 PTPRC HDAC9 MAFB HSPH1 SAMSN1 NFKBIZ LYN
TNFA signaling via NFKB
Lymphocyte activation

**Supplemental figure 8. Developmental changes in gene expression in the fetal lung immune cells**
a) pathways induced during fetal lung development (14+1 until 19+0 weeks) in the immune cells as identified by enrichment analysis based on the DSA. b) Heatmaps depicting the enriched terms associated with individual GAs as identified by multi-list enrichment analysis based on DSA in immune cells. c) Heatmap depicting the expression level of genes associated with selected enriched terms as identified in panel b). Expression levels in the heatmaps and dotplots are presented as log(TP10k+1) values. Log(TP10k+1) corresponds to log-transformed UMIs per 10k.
